# Bound or unbound: Mapping and monitoring receptor oligomerization using time-resolved fluorescence

**DOI:** 10.64898/2026.02.21.707147

**Authors:** Annemarie Greife, Ruiqi Liu, Paul S. Koehler, Katrin G. Heinze, Katherina Hemmen, Thomas-Otavio Peulen

## Abstract

Understanding protein oligomerization in living cells is essential for elucidating cellular signaling and regulation, yet quantitative analysis remains challenging due to heterogeneous expression levels, dynamic interactions, and limited access to absolute protein concentrations. Here, we present a standardized, open-source framework for quantifying protein assemblies in living cells by integrating fluorescence lifetime and anisotropy imaging (heteroFRET and homoFRET) with molecular brightness-based concentration estimation and image analysis.

Using natural variants of a vertebrate GPCR, the melanocortin-4 receptor (MC4R-A and MC4R-B2), as a model system, we demonstrate how to discriminate monomers, dimers, and higher-order oligomers, extract inter-fluorophore distance distributions, and determine association constants under physiologically relevant conditions in living cells. Standard fluorescent protein tags report on proximity and oligomerization via Homo- and HeteroFRET. Association constants are quantified using the variable protein expression in living cells and the spectroscopy readouts. By high-content imaging we overcome the biological noise and attain data qualities comparable to conventional biochemical *in vitro* assays. Intensity- and fluctuation-based segmentation further extends the accessible concentration range within individual cells, improving affinity analysis robustness.

Our results establish quantitative image spectroscopy on living cells as quantitative tool for investigating protein-protein interactions under physiologically relevant conditions. All computational workflows are implemented in open-source software and are accompanied by detailed protocols and analysis scripts, enabling reproducible application and adaptation. Beyond GPCRs, this framework provides a practical and transferable methodology for quantitative studies on protein-protein interactions, mechanistic studies and drug discovery in complex cellular environments.

## 1 Introduction

Proteins orchestrate nearly every biological function. In many signaling processes in and between cells, individual proteins assemble into protein complexes. Thus, it is essential to understand the composition and interaction dynamics of such protein complexes to better disclose their biological and physiological roles (Marsh and Teichmann 2015). Classical approaches such as yeast two-hybrid (Duarte and Euclydes 2024), co-immunoprecipitation (Tan and Yammani 2022), and pull-down assays (Gnanasekaran and Pappu 2023) provide highly valuable information on protein-protein interactions (PPI). These classic approaches are efficient and successful for soluble cytoplasmic or nuclear proteins, but can be challenging for transient interactions, and membrane-bound proteins (Zhang et al. 2025). Here, Förster Resonance Energy Transfer (FRET) (Förster 1948) offers a clear advantage, as interactions in living cells can be spatially monitored in real-time (Bonilla and Shrestha 2024, Sun et al. 2012, Zhang, et al. 2025).

Among all membrane-bound proteins, the largest family are the G protein-coupled receptors (GPCR), which are important pharmacological drug targets. About 35% of approved drugs so far target GPCRs or GPCR-related proteins (Calebiro and Grimes 2020, Sriram and Insel 2018). Melanocortin 4 receptor (MC4R) is a class A GPCR involved in metabolism and obesity in humans (Farooqi et al. 2003). MC4R mainly expresses in the central nervous system, particularly in the hypothalamus (Wei et al. 2023). It plays a key role in energy homeostasis, including both food intake and energy expenditure. The MC4R system is highly conserved across vertebrates, from fish to mammals (Tao 2022). In *Xiphophorus* fish, commonly known as platy fish and swordtails, the highly polymorphic and sex-linked *mc4r* locus is the key genetic factor controlling the timing of puberty and, as a result, the ultimate body size in males (Lampert et al. 2010, Volff et al. 2013). This leads to a remarkable divergent expression profile within their natural population. In two swordtails, *Xiphophorus nigrensis* and *Xiphophorus multilineatus*, onset of puberty is attributed to the ratio of the *mc4r* alleles A, B1, and B2, and their copy number variation on the X- and Y-chromosomes (Lampert, et al. 2010). The wild-type A allele has full signaling capacity, while the mutant B alleles, B1 and B2, cannot be stimulated by agonists. Unlike A allele, B1 and B2 lack the C-terminal dicysteine motif and unable to anchor the C-terminal helix VIII into the plasma membrane. B2 has another four-base deletion, resulting in an elongated C-terminus (Lampert, et al. 2010). Our previous studies on MC4R receptor dimerization revealed that wildtype MC4R-A receptors homo- and hetero-dimerize. Homomers are fully active, in contrast to MC4R-B receptors. MC4R-B1 did not self-associate, but formed a signaling incompetent hetero-dimer with MC4R-A. MC4R-B2 formed stable homo-and hetero-dimers, thereby exerting a dose-dependent dominant-negative effect on ligand binding and signal transduction (Liu et al. 2020, Liu et al. 2023).

It has been recognized that class C GPCRs are fully functioning only as dimers to transduce signals (Kniazeff et al. 2011); yet for class A GPCRs, such as MC4R, debate remains if and to which extent they form dimers or oligomers, and what function the dimer/oligomer confers (Milligan et al. 2019). *In vitro* analyses only provide limited information about interaction in living cells (Zhang, et al. 2025). To study PPIs in living cells, we combine fluorescence spectroscopy with high-content imaging and establish open-source analysis workflows for quantitative PPI analysis (Peulen 2025, Peulen et al. 2025), and provide step-by-step protocols (Greife et al. 2025). As benchmarks to investigate oligomerization states, we use two *Xiphophorus* MC4R variants, A and B2, tagged with standard fluorescent proteins (FPs) at the C-terminus, representing receptors with respective short and long peptide chains linking to the FPs (Lampert, et al. 2010, Liu, et al. 2023). With this receptor model, we apply time-resolved FRET to measure GPCR proximity via Homo- and Hetero-FRET.

The non-radiative energy transfer from a donor to an acceptor fluorophore (Förster resonance energy transfer, FRET) can report on inter-fluorophore distances (Förster 1948). The efficiency of that transfer process, *E*, depends on the sixth power of the inter-fluorophore distance (*R_DA_*). Thus, FRET happens typically at distances below 10 nm. As a molecular ruler, FRET is uniquely suited to report molecular proximity and conformational or interaction changes in living cells (Sun, et al. 2012, Weidtkamp-Peters et al. 2009, Zhang, et al. 2025). Classical intensity-based FRET approaches often suffer from spectral bleed-through, cross-excitation, and variable fluorophore expression levels (Bonilla and Shrestha 2024). Time-resolved, fluorescence lifetime-based FRET implemented through time-correlated single photon counting (TCSPC) or frequency-domain fluorescence lifetime imaging (FLIM) mitigate these limitations by directly measuring FRET-induced donor lifetime shortening (Bonilla and Shrestha 2024, Weidtkamp-Peters, et al. 2009). FLIM-based FRET is largely independent of fluorophore concentration, photobleaching, or excitation intensity and therefore provides a quantitative, calibrated approach to probe interaction dynamics in heterogeneous samples such as the plasma membrane (Greife et al. 2016), cellular compartments (Kravets et al. 2016), or in crowded cellular microdomains (Lou et al. 2024).

Depending on fluorophore configuration, two types of FRET experiments inform on PPIs. In heteroFRET experiments, distinct donor and acceptor fluorophores inform on pairwise interactions. They are particularly useful for mapping binary associations and the fraction of interacting molecules (Greife, et al. 2016, Kravets, et al. 2016, Weidtkamp-Peters, et al. 2009, Zhang, et al. 2025). In homoFRET experiments, the energy transfer between identical fluorophores is monitored. The energy migration between two identical fluorophores manifests as a decrease of fluorescence anisotropy. HomoFRET experiments allows single-color detection of oligomer states (Chan et al. 2011, Warren et al. 2015) and are therefore suited to quantify concentration-dependent oligomerization or to distinguish monomers from dimers/oligomers (Chan, et al. 2011). HomoFRET experiments are technically and analysis-wise more demanding, because the time-resolved fluorescence anisotropy decays are influenced by rotational diffusion, local viscosity, and photoselection; consequently, robust decay models including the required corrections and appropriate instrument corrections are essential to extract reliable donor-donor energy transfer rates (Vogel et al. 2015, Vogel et al. 2009).

Until recently, the implementation of quantitative FRET/FLIM - especially time-resolved anisotropy for homoFRET - required proprietary software or scripting, and substantial expert knowledge. This limited a broad adoption, particularly when access to specialized spectroscopy microscopes and workflows is also limited. Over the last years, substantial progress has been made in open-source software ecosystems that integrate photon-counting, lifetime fitting, anisotropy decay modeling, ROI-based quantification, and spatially resolved FRET efficiency mapping. Packages such as FLIMfit (Warren et al. 2013), PAM (Schrimpf et al. 2018), ChiSurf (Peulen 2025), and modular Python-based workflows, e.g. in *tttrlib* (Peulen, et al. 2025), now allow non-experts to perform data analysis and complex model fitting, correct for scattered light and objective depolarization, use model-free estimators such as *ε*(t), and incorporate segmentation or concentration maps directly into pixel-wise analysis. These developments enable reproducible, transparent oligomerization studies in live cells, lower the entry barrier for quantitative FRET/FLIM, and provide a foundation for standardized protocols. In this work, we build upon these recent advances and provide practical guidelines, quality checkpoints, and experimental considerations for reliable FRET-based oligomerization analysis using MC4R receptor variants as GPCR model system.

Conceptually, this work establishes a practical and quantitative framework for analyzing protein oligomerization in living cells by integrating time-resolved heteroFRET and homoFRET, molecular brightness-based concentration estimation, and image segmentation. By combining these complementary modalities within an open-source analysis workflow, we overcome technical hurdles in high content image spectroscopy and key limitations of conventional FRET approaches, including uncertainty in protein concentration, heterogeneous expression, and ambiguity in oligomeric state assignment. Our approach enables the direct extraction of apparent association constants and inter-fluorophore distance distributions under physiologically relevant conditions, while maintaining experimental and computational accessibility. Beyond the specific case of GPCRs, this framework provides a methodology for quantitative studies of protein-protein interactions in cells.

## 2 Material and Methods

### 2.1 Sample preparation & Data acquisition

#### 2.1.1 Plasmid preparation

The genes for *mc4r-A* and *B2* have been cloned previously into a pcDNA3.1 backbone between *HindIII* and *XbaI* while eGFP and mCherry, respectively, were cloned C-terminally using *XbaI* and *NotI* restriction sites (Lampert, et al. 2010, Liu, et al. 2023). No further amino acids were inserted as linkers between MC4R and the fluorescent protein (**Supp. Table 1**).

#### 2.1.2 Cell culture

Human embryonic kidney 293T (HEK293T) cells were cultured in Dulbecco’s modified Eagle medium (DMEM) without sodium pyruvate (P04-03550, PAN Biotech, Aidenbach, Germany), with 10% fetal calf serum (AC-SM-0190, Anprotec, Bruckberg, Germany) and 100 U/mL penicillin, 100 µg/mL streptomycin (P4333, Sigma-Aldrich, St. Louis, Missouri, United States) at 37°C, 5% CO_2_. To passage the cells, culture medium was aspirated; cells were washed with PBS (14190, Gibco, Thermo Fisher Scientific, Waltham, Massachusetts, United States) once, and cells were detached using 0.5× Trypsin-EDTA solution (T4299, Sigma-Aldrich). The cell suspensions were resuspended in DMEM culture medium. Cells were regularly passaged every 3-4 days.

#### 2.1.3 Live-cell Fluorescence Lifetime Imaging (FLIM)

FLIM was performed on live-cell samples. HEK293T cells were seeded at 75’000 cells/well on poly-D-lysine coated 4-chambered coverglasses (C4-1.5P, Cellvis, Mountain View, California, United States) and grown for ∼48 h. Cells were transiently transfected by jetPRIME using 1 µL jetPRIME reagent and 50 µL jetPRIME buffer (Sartorius Sartorius GmbH & Co. KG, Goettingen, Germany) according to the manufacturer’s instructions with either an eGFP-tagged construct for DO samples or a mixture of eGFP- and mCherry-tagged constructs for the FRET samples. In each sample, a total of 0.5 µg DNA was used in a transfection volume of 500 µL DMEM. Empty backbone vector was mixed with MC4R constructs to keep the total DNA amount constant. For DO samples, 0.25 µg eGFP-tagged constructs were mixed with 0.25 µg empty pcDNA3 vector. For FRET samples, the ratio between eGFP- and mCherry-tagged construct varied from 1:1 to 1:20. The cell culture medium was changed at six to eight hours post-transfection, and the cells were incubated overnight. On the next day, medium was changed to imaging medium [DMEM without phenol red (P04-01161, PAN Biotech) supplemented with 15 mM HEPES (15630-080, Gibco), 10% fetal calf serum, and 2 mM glutamine (G7513, Sigma-Aldrich)], which allows for 2-3 h of live cell imaging without a CO_2_ source.

##### Setup

FLIM was performed using a Zeiss LSM980 confocal microscope (Zeiss, Oberkochen, Germany), operated with ZEN3.3 software and equipped with the LSM upgrade kit (PicoQuant, Berlin, Germany), which is managed by SymPhoTime64 software. Both systems are controlled through the Zeiss Blue PicoQuant Application software. Live-cell samples were imaged using a 40× 1.2 NA water objective. The excitation was achieved using pulsed lasers at 485 nm and 560 nm (LDH-P-C-485B/LDH-D-TA-560B, PicoQuant, Berlin, Germany). The emitted fluorescence photons were first split by a polarizing beam splitter, followed by a 560 nm long pass (560/LPXR, AHF, Tübingen, Germany) to separate green and red signals. Green photons passing HC520/35 bandpass filters (Semrock, New York, U.S.A.) were registered by PMA Hybrid 40 detectors (PicoQuant, Berlin, Germany). Red photons passing an ET600/50 bandpass filter (Chroma Technology GmbH, Olching, Germany) were detected by Excelitas SPADs (SPCM-AQRH-14-TR, PicoQuant, Berlin, Germany).

##### Image acquisition

Images were acquired at a zoom factor of 6.0 resulting in a pixel size of 0.069 µm. The image dimensions were set at 512 × 512 pixels, and the data was collected at a pixel dwell time of 8.16 µs over the course of 50 frames.

##### FLIM measurement

During the measurement, the repetition rate was set to 40 MHz. Pulsed interleaved excitation (PIE) (Muller et al. 2005) was utilized at a delay time of 25 ns and for a time window of 50 ns to modulate donor and acceptor excitation. The excitation laser power was adjusted between 200 and 800 nW at 480 nm and 140 and 550 nW at 560 nm. The time-correlated single photon counting (TCSPC) resolution was 10 ps.

##### Calibration procedure

Before live-cell measurements, the setup was calibrated following a standard routine. Initially, the instrument response function (IRF) was determined using a saturated solution of Erythrosine B (200964, Sigma-Aldrich) in potassium iodide (KI) (221945, Sigma-Aldrich). Daily fluorescence correlation spectroscopy (FCS) and TCSPC data acquired for the dyes Alexa488 and Alexa568 and the fluorescent proteins eGFP and mCherry served as molecular brightness and diffusion standards to determine molecular concentrations, the instrumental g-factor and polarization mixing factors (Erdelyi et al. 2014, Koshioka et al. 1995).

#### 2.1.4 Simulation of fluorescent protein distributions

The sterically allowed distribution of fluorescent proteins, FPs, coupled to MC4R dimers were simulated using the integrative modelling platform, IMP (Russel et al. 2012). Given the amino acid sequences, dimeric structures of MC4R with attached FPs were created with AlphaFold3 (Abramson et al. 2024). To effectively sample the sterically allowed conformational space, the structures were coarse-grained in IMP on residue-level. Residues and groups of residues are represented by beads. In the coarse-grained representation MC4R dimers and FPs were represented by rigid bodies, whereas the linker coupling the FPs to the MC4R dimer was represented by a string of flexible beads. In rigid bodies, relative distances across beads are constrained during conformational sampling. In flexible strings of beads, the beads are restrained by the sequence connectivity (Algret et al. 2014). In the sampling, excluded volume restraints (Shi et al. 2014) were applied in addition to a knowledge-based membrane restraint localizing the MC4R dimer into a membrane and excluded the FPs and the linker from the membrane. The sterically allowed models were sampled by Gibbs sampling, based on the Metropolis Monte Carlo algorithm. FP, MC4R, and bead positions were randomized followed by a brief gradient descent optimization. Next, Monte Carlo moves included random translation and rotation of rigid bodies (up to 1 Å and 0.01 radians, respectively), and random translation of individual beads in the flexible segments (up to 2 Å) were used for sampling. For each computed model, the inter FP distance, *R*_*DA*_, and the orientation factor, ***κ***^2^, were calculated.

### 2.2 Data analysis

#### 2.2.1 Live-cell Fluorescence Imaging Imaging (FLIM)

Our time-resolved FRET analysis workflow consists of several steps (outlined in detail below). Briefly, the registered photons were binned into pixels to yield images. Registered photons encode information on detectors and their arrival time with picosecond resolution. We compute pixel-wise intensities (intensity images) and mean photon arrival times relative to the excitation pulse as average features that we map to images. These features we use to identify cells, segment images, and to define regions of interest (ROIs). Finally, we integrate photons of ROIs into fluorescence intensity decay histograms (fluorescence decay). Integrating photons of ROIs decreases the noise of fluorescence decays and enables fitting of the fluorescence lifetimes and inter-fluorophore distances.

##### Export of intensity images

Fluorescence intensity images were generated by integrating photons of all frames in the three specified channels: green-prompt (donor emission), red-prompt (FRET-sensitized acceptor emission), and red-delay (directly excited acceptor emission). The data export into 16-bit tiff-files was programmed in scripts using *tttrlib* (Peulen, et al. 2025) and *scikit-image* (van der Walt et al. 2014).

##### Image segmentation

To define cells and regions of interest (ROIs) for each acquired image, the total intensity sum of all three channels (green prompt, red prompt and red delay) was used. The image of the intensity sum was in a first step smoothed by using a median filter with a two-pixel radius, followed by Li thresholding as implemented in *scikit-image* (Li and Lee 1993, Li and Tam 1998). In the next step, holes smaller than 200 pixels were filled and small objects (e.g. cells at the image border) encompassing less than 10’000 pixels removed. The binary images, *i.e.* the cell masks, were exported as 8-bit tiff files with intensity values = 0 inside the cells and 255 outside the cell. The automated segmentations were controlled and curated in Fiji (Schindelin et al. 2012).

##### Sub-region segmentation

Within the cell membrane the fluorescence intensity was inhomogeneous. To assess if intensity variations and dimerization differences are related, ROIs were segmented into five different sub-ROIs using two distinct methods. <u>In the first method</u>, the vesicles detected mainly in the red channels were framewise selected in the red delay time series using an *ilastik* based segmentation (Berg et al. 2019). In *ilastik*, a pixel classifier is trained based on user input to recognize structures of interest. We used the cell density counting approach implemented in *ilastik* trained on manually annotated examples from representative images to generate pixel-wise vesicle probability maps (sigma=2.50) and exported binary cell masks. In the next step, the green prompt time series were loaded jointly with the vesicles mask and the cell mask. All pixels belonging to the vesicles were removed from the cell mask and the obtained cell mask was applied onto the green prompt intensity images to build frame integrated intensity images (sum image). The sum image intensity histogram was used to split cells into a “low” and a “high” intensity region by Otsu’s thresholding implemented in *scikit-image* (Otsu 1979). <u>In the second method</u>, we used the “Number & Brightness” (N&B) approach (Digman et al. 2008), that relies on the intensity fluctuations along the image time series to elucidate an pixel-wise brightness and apparent number of molecules. We compute intensity fluctuations based on the total intensity in all channels (green prompt, red prompt and red delay) and split cells in a “low B” (brightness below 1.375, assumed to be rather monomeric) and a “high B” brightness region (brightness above 1.375, assumed to be rather dimeric). The segmented five different regions were converted into binary masks, saved as 8-bit tiffs, and used to export the fluorescence intensity decays and other properties of these regions.

##### Export of fluorescence intensity decays

The exported binary ROI images from the image segmentation were used as selective mask to compute the fluorescence decay histograms of the ROI using *tttrlib* and *scikit-image*. The fluorescence decay histograms were computed for all channels (green prompt, red prompt and red delay) with a twofold micro time binning resulting in a histogram bin size of 20 ps. The signal from the parallel and perpendicular detection channel was stacked into a single column.

##### Determination of correction factors

FLIM measurements were performed with vertical polarized excitation (V) and detection, either in plane (VV) or 90° rotated (VH) to the excitation direction. The used high NA objectives lead to polarization mixing that needs to be considered to compute artifact free fluorescence intensity decays (Erdelyi, et al. 2014, Koshioka, et al. 1995). Moreover, the detection sensitivities of the parallel and perpendicular detectors need to be corrected by the g-factor, *g*, typically defined as the sensitivity ratio of the parallel (*I*_*VV*_) and perpendicular (*I*_*VH*_) detector. In an ideal system, *g* is close to 1. A g-factor of two means that the detection system is twice as sensitive to vertically polarized light as it is to horizontally polarized light. To determine *g*, *l_1_*, and *l_2_* we jointly analyze the fast rotating fluorophores Alexa488 and Alexa568 along with slowly rotating fluorescent proteins (FPs) in *ChiSurf* (Peulen 2025) using a single rotational correlation time and mono- (Alexa dyes) or bi-exponential fluorescence lifetime models (FPs). Details on the fitting procedure can be found in the software manual. On average, the green g-factor was 0.946 with *l_1_* and *l_2_* being 0.059 and 0.305, respectively. The red g-factor was 0.995, and *l_1_* and *l_2_* were 0.121 and 0.279, respectively.

##### Determination of average fluorescence lifetimes

Average fluorescence lifetimes of eGFP in eGFP transfected (Donly/DO) and in eGFP and mCherry co-transfected FRET samples (DA) for the eGFP fluorescence channel (488 nm excitation) were determined in *ChiSurf* by fitting by iterative reconvolution of multi-exponential models to the exported time-resolved fluorescence intensities:

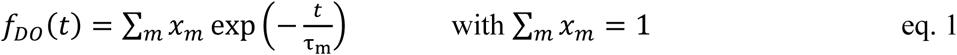

Briefly, experimental nuisances were introduced into the model by convolutions with the IRF, computing the partial polarization mixing, and scaling models by the detection sensitivity of the detectors. For a model, the average fluorescence lifetime 〈*τ*〉_*x*_ was calculated as species-average:

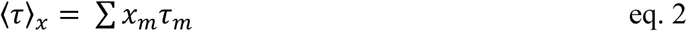

##### Determination of inter-fluorophore distances (heteroFRET)

Inter-fluorophore distance distributions in FRET samples were determined by fitting a mixture model *f*(*t*) to the data, which contains the fluorescence decay of non-FRET molecules, *f*_*DO*_(*t*), and *f*_*DA*_(*t*) = *∈*_*D*_(*t*) ⋅ *f*_*DO*_(*t*), the fluorescence decay of FRET species:

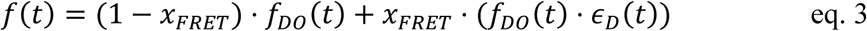

Here, *x*_*FRET*_ is the fraction of FRET active species and *∈*_*D*_(*t*) is the FRET-induced donor decay that can be directly related to FRET rate constants and distances (Peulen et al. 2017).

Due to the flexible C-terminal tagging of the FPs, we relate the FRET-induced donor decay to a distance distribution. In this analysis, use a Gaussian distribution of half-width *σ*_*app*_and centered around *R̄*_*app*_. In this analysis, we compute FRET-rate constants for distances for a single averaged ***κ***^2^ of 2/3, yielding apparent inter-fluorophore distances; potential deviations due to restricted fluorophore orientations are addressed in the **Discussion**. Accordingly, the recovered distances represent apparent distances, *R*_*app*_, that reflect both spatial separation and orientational averaging rather than absolute fluorophore separation. Nevertheless, these recovered distances directly relate to the FRET-induced donor decay *∈*_*D*_(*t*) by:

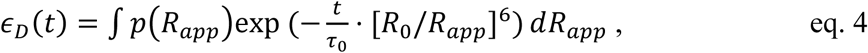

where the inter-fluorophore distribution *p*(*R*_*app*_) is approximated by a normal distribution:

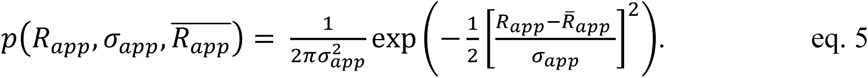

Here, we used a literature value of 52 Å for the used eGFP/mCherry FRET-pairs (Lambert 2019).

<u>In the first analysis round</u>, we fitted the fluorescence decays of entire cells by a single Gaussian *R̄*_*app*_and *σ*_*app*_ (limited to 6 Å). <u>In the second analysis round</u>, we fitted the fluorescence decays of entire cells by a mixture of two Gaussians. The longer distance, *R̄*_*di*_represents the MC4R dimer distance (*σ*_*di*_ranged from 5 to 25 Å). In the presence of multiple acceptors, all acceptors contribute to FRET (sum of quenching processes). Accordingly, a short apparent distance with a narrow distribution (*σ*_*app*_ ≈1 Å) was used as an effective description of collective quenching by multiple acceptors in oligomeric assemblies (Greife, et al. 2016, Kravets, et al. 2016). <u>In the third analysis round</u>, the average values for *R̄*_*di*_, *R̄*_*ol*_and *σ*_*di*_ were fixed to determine species fractions for the entire cells and the five segmented cell ROIs.

##### Inter-fluorophore distances determined by homoFRET

Fluorophores show fluorescence depolarization due to their rotational motion. FPs alone have a rotational correlation time, *ρ*_*FP*_, of around 12 ns (Striker et al. 1999). For the FP coupled to MC4R we estimated that the rotational correlation is slower (*ρ*_*global*_ ≈ of 100 ns). In homoFRET, a FRET-induced (additional) fluorescence depolarization of the emitted light reports on fluorophore proximity. In homoFRET the relaxation time, *ρ*_*FRET*_, decreases with the distance, eventually lowering the initial amplitude of the fundamental anisotropy, *r*_0_. We jointly fitted the parallel, *I*_*VV*_(*t*), and perpendicular, *I*_*VH*_(*t*), fluorescence decays using bi-exponential (dimer) and tri-exponential models (oligomer):

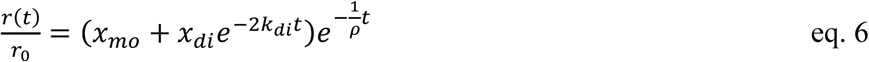

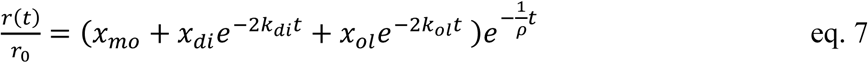

Here, *ρ* is the global rotational correlation time of the protein bound FP, *x*_*mo*_, *x*_*di*_, and *x*_*ol*_ are species fractions of the monomer, dimer, and oligomer, respectively, with corresponding FRET-rate constants *k*_*di*_ (dimer) and *k*_*ol*_ (oligomer).

In our data analysis, we described *r*(*t*) by a multiexponential decay and determine relaxation times *ρ*_*di*_ and *ρ*_*ol*_, which are coupled to the global molecule rotation, *ρ*:

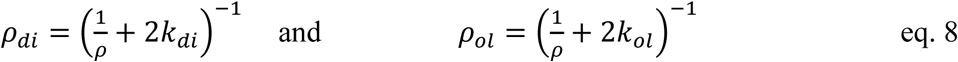

Note, in contrast to unidirectional heteroFRET, a factor of two is included in the equation as homoFRET occurs bidirectional between donor molecules. We relate the FRET-rate constants to apparent distances in the dimer, *R*_*di*_, and oligomer, *R*_*ol*_, using the Förster equation:

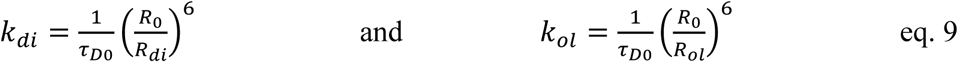

To obtain *R*_*di*_ and *R*_*ol*_ for measured relaxation times *ρ*_*di*_ and *ρ*_*ol*_:

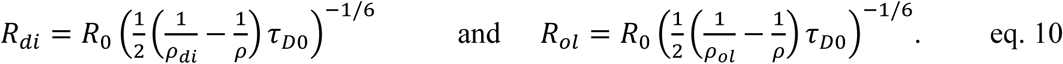

Here, we used *R*_0_= 49 Å and *τ*_*D*_= 2.38 ns for eGFP and *R*_0_= 44 Å and *τ*_*D*_= 1.31 ns for mCherry (Lambert 2019). Note, analogous to heteroFRET samples, distance distributions are a more accurate description. Nevertheless, for simplicity in the analysis, we fitted discrete relaxation times, *ρ*, and computed corresponding FRET-rate constants.

In addition to the time-resolved anisotropies, we compute the steady-state anisotropy, *r*_*ss*_, of eGFP in eGFP-only samples, eGFP/mCherry transfected samples and *r*_*ss*_ of directly excited mCherry using the background-corrected count rates in the parallel, *S*^*BG*^, and perpendicular channels, *S*^*BG*^.

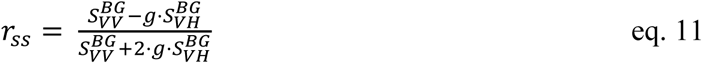

Here, the background of the channels was estimated by the fitted offset of the fluorescence decay curves in the respective detection channel.

#### 2.2.2 Concentration estimation: Converting photon counts to concentration

To relate the changes in dimer or oligomer fractions to the MC4R concentration, we performed fluorescence correlation spectroscopy (FCS) on reference samples in singly transfected (MC4R-eGFP or MC4R-mCherry) cells to determine the molecular brightness (counts per molecule per second, cpms) of the fluorescent proteins. Using these experimental cpms, the measured fluorescence intensities can be converted to the number of molecules and concentrations (Hemmen et al. 2021). To determine concentration, the detection volume needs to be known. Using Alexa488 and Alexa568 as reference, we determined average detection volumes of ∼ 0.81 fl (488 nm) and 0.92 fl (560 nm), respectively. At an excitation intensity of 0.54 µW, we obtained a molecular brightness of 92 cpms for eGFP and 37 cpms at 0.3 µW for mCherry. The concentration of mCherry, [A], was determined based on the background-corrected average count rate, in a ROI:

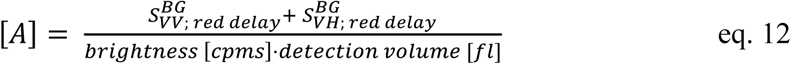

For the eGFP concentration in the donor only cells, [D_0_], the concentration estimation was performed similarly:

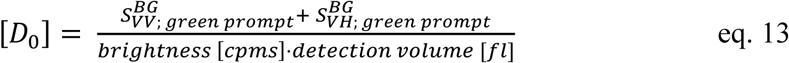

For the eGFP concentration in the FRET samples, the background-corrected average count rate was corrected for the quenching due to FRET:

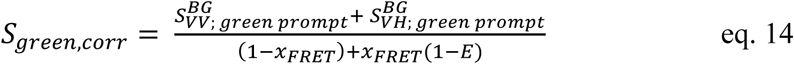

The energy transfer efficiency *E* and the fraction of the FRET-active population, *x_FRET_*, were taken from the Gaussian distance fit described above (**eq. 3-5**). Subsequently, the concentration of eGFP in the presence of mCherry [*D*_*A*_] was estimated using the corrected count rate:

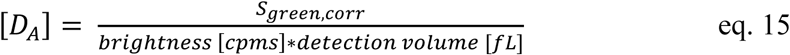

The ratio of the donor to the acceptor (or total protein, *T*) concentration influences strongly the observed amount of FRET. Thus, we define the ratio as

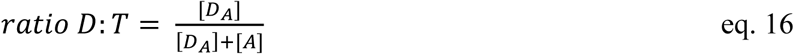

#### 2.2.3 Estimation of the association constants for oligomerization

To describe the dimerization and oligomerization of the MC4R variants, a simple stepwise model is used; in a pre-equilibrium the dimer is formed first, followed by subsequent oligomerization, similar to what was described previously (Greife, et al. 2016, Kravets, et al. 2016). In this analysis higher-order oligomers are formally represented by a tetrameric species.

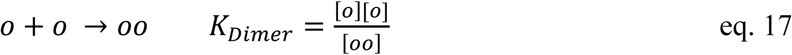

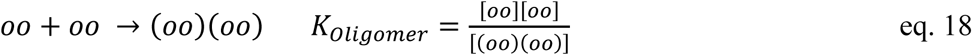

Here o is a monomer, oo a dimer and (oo)(oo) is a tetramer. We use the monomer o as the base species. Then the total protein concentration is given by:

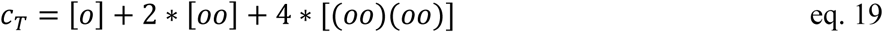

Now, the three species fractions *x_mo_*, *x_di_* and *x_ol_* for any given total protein concentration are used to determine *K*_*Dimer*_ and *K*_*Oligomer*_ by solving the three equations above.

#### 2.2.4 Estimation of the dimerization constant from homoFRET analysis

For the estimation of an apparent dimerization constant, *K*_*Dimer*,*app*_, we merged the fit results from the DO and directly excited acceptor anisotropy analysis. While the acceptor signal was analysed with the dimer model (**eq. 6**), the DO samples were best described with an oligomerization model (**eq. 7**). Thus, for the DO samples we combined the species fractions *x*_*Di*_ and *x*_*Oligo*_to yield *x*_*Di*,*app*_. Similarly, as above, we define *K*_*Dimer*,*app*_ and *c*_*T*_, and solve the two equations:

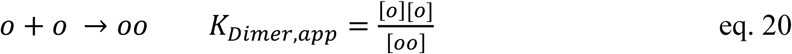

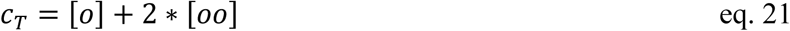

##### Pixel-based mean photon arrival times

The donor mean photon arrival time was calculated from the micro time distribution for the stacked frames using *tttrlib* and *scikit-image*. Only pixels with more than 15 detected photons per frame were considered and the result was exported as 32-bit float tiff-image. To visualize only a selected ROI, the binary ROI mask from the segmentation step was multiplied with the mean arrival time image.

#### 2.2.5 Statistical analysis

Statistical analyses were performed in OriginPro 2021b; (OriginLab Corporation, Northampton, MA, United States) using ANOVA. * = *p* < 0.05, ** = *p* < 0.01, *** = *p* < 0.001; individual cells were treated as independent observations, and no additional correction for experimental nesting was applied.

#### 2.2.6 Excluded data

All data points underwent a 2-dimensional, multivariate location and scatter analysis. To detect outliers, the Mahalonobis distance was calculated (Etherington 2021). Samples were excluded using a 97.5% threshold of the Mahalonobis distance determined from the monomer, dimer, and oligomer species fractions of the final global fit and the donor to total protein ratio.

## 3 Results

### 3.1 FRET as a tool to identify protein-protein interactions

Wild-type MC4R receptors can form both homodimers, composed of two wild-type receptors, and heterodimers with mutant variants. This dimerization modulates intracellular signaling strength and may contribute to the variations observed in the timing of puberty (Lampert, et al. 2010, Liu, et al. 2020, Liu, et al. 2023). In this study, we build on the established knowledge of MC4R homodimerization (Kleinau et al. 2020, Liu, et al. 2023, Nickolls and Maki 2006, Piechowski et al. 2013) to develop a detailed analysis workflow for live cell experiments to (*i*) detect dimers and potential higher-order oligomers in live cells, (*ii*) estimate of association constants (*K_D_*), (*iii*) enhance discrimination between monomeric, dimeric, and oligomeric species using spectroscopic and image-derived features, and (*iv*) construct structural and geometric models.

All constructs in this study are C-terminally fused to fluorescent proteins, either eGFP or mCherry (**Fig. 1A**). The wild-type MC4R-A possesses a comparatively short C-terminal tail that includes a post-translationally modified di-cysteine motif which anchors the cytoplasmic helix VIII to the cell membrane (Lampert, et al. 2010, Liu, et al. 2023). In contrast, the B2 variant contains a frameshift mutation that removes this motif and results in an extended C-terminal tail. Consequently, MC4R-A and -B2 differ in their flexible amino acid sequence (linker length), possibly altering the orientational (*κ*^2^) and spatial freedom (accessible volume) of the attached fluorophores at the membrane.

**Figure 1.**
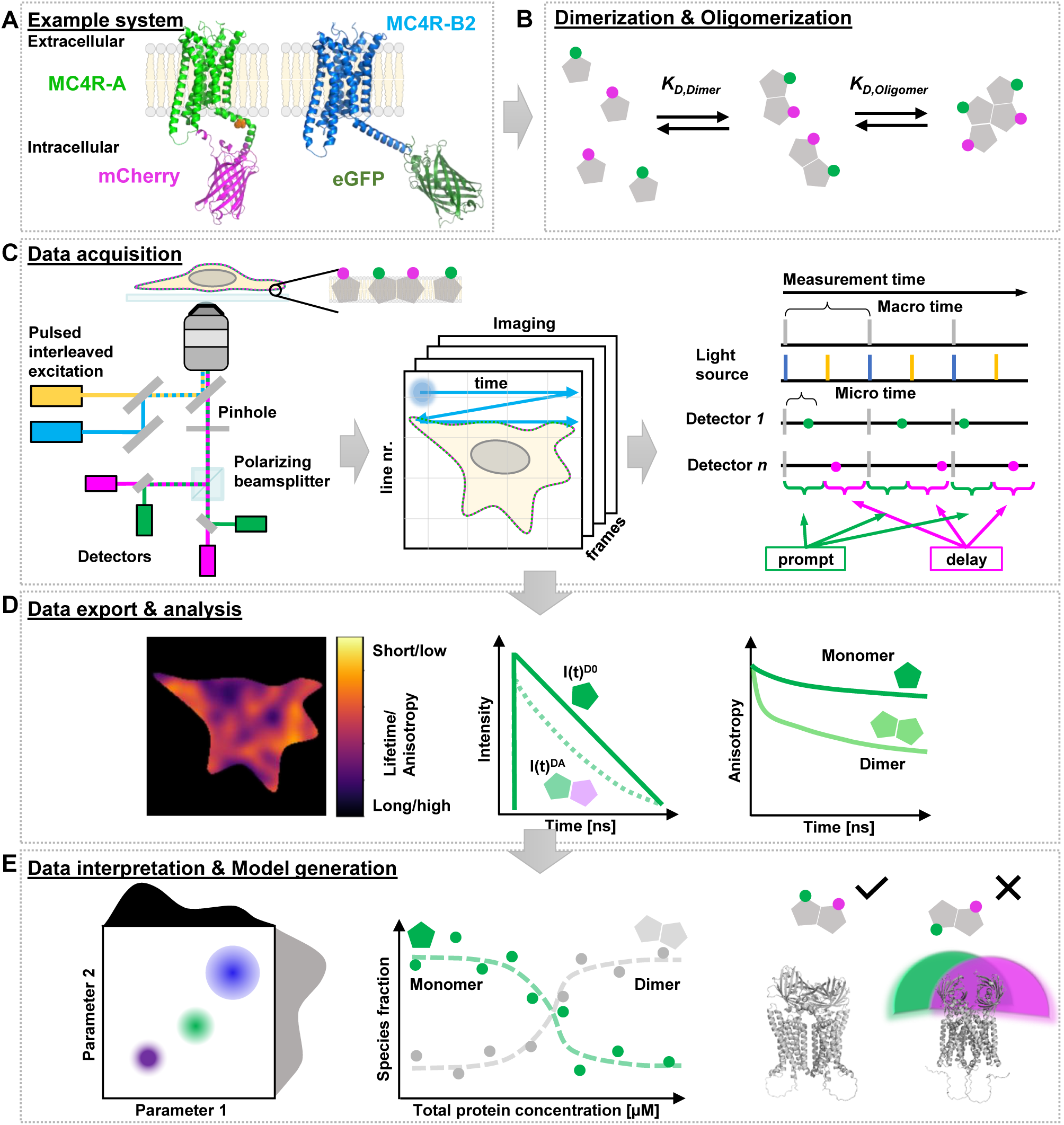
Fluorescence Lifetime Imaging (FLIM) combined with Pulsed Interleaved Excitation (PIE) resolves protein-protein interaction in the melanocortin-4 receptor (MC4R) by homo- or heteroFRET. **(A)** We focus on two out of three alleles from *Xiphophorus* MC4R, which differ in their intracellular C-terminal domain. In the A type (light green), two membrane anchors (orange) hold the intracellular C-terminal domain close to the membrane. In the B2 variant (blue), a frameshift causes the loss of the two membrane anchors, resulting in a much longer C-terminal tail. The MC4R structures were generated using AlphaFold3 (Abramson, et al. 2024). **(B)** C-terminally tagged constructs with either mCherry (magenta) or eGFP (green) (simplified as pentagons with magenta or green circles) are used for affinity and multimerization analysis: monomers are represented as pentagons, dimers and higher order oligomers are represented as connected pentagons. **(C)** In confocal polarization-resolved PIE-FLIM, the donor (green, 485 nm laser) and the acceptor (magenta, 561 nm laser) excitation are alternated while the image is scanned. The photons’ arrival times relative to the experiment start (macro time) and the preceding laser pulse (micro time) are recorded, along with the detection channel (green/magenta, parallel/perpendicular) and image frame markers. **(D)** The information encoded in the photons is used to construct mean arrival times and steady-state anisotropy images <u>(left</u>). The integrated micro time of images or region of interest (ROIs) is analyzed to determine fluorescence lifetimes, inter-fluorophore distances (<u>middle</u>), or time-resolved anisotropies (<u>right</u>). **(E)** Joint analysis of multiple parameters and spatial information allows to disentangle subcellular features (<u>left</u>). Exploiting varying expression levels allows for probing oligomerization and association constants (<u>middle</u>). Structural models of proteins (grey) and fluorophores (green/magenta) allow extraction of relative protein orientations and dimerization interfaces (<u>right</u>).

Our analysis focuses on the homodimerization and potential higher-order oligomer formation of MC4R-A and MC4R-B2 (**Fig. 1B**). To characterize these interactions, we performed fluorescence lifetime imaging (FLIM) in pulsed interleaved excitation (PIE) mode for both receptor variants (Muller, et al. 2005, Weidtkamp-Peters, et al. 2009). In live-cell PIE-FLIM, eGFP (donor) and mCherry (acceptor) are excited alternately on the nanosecond timescale: the donor is excited in the prompt time window, followed by direct acceptor excitation in the delayed time window (**Fig. 1C**). Emitted fluorescence is collected through the same objective and separated first by polarization (vertical, VV, or horizontal, VH, relative to the excitation direction) and then by the emission color (donor vs. acceptor).

During data acquisition, the laser scans the image across multiple frames. For each detected photon, several parameters are recorded: (*i*) the detector identity (green/red and polarization channel), (*ii*) the macro time (time since acquisition started), and (*iii*) the micro time (time since the last prompt laser pulse). The macro time, together with encoded line and frame markers, allows assignment of photons to individual pixels. The micro time distinguishes whether a red photon originates from FRET-sensitized emission - indicating donor excitation followed by energy transfer - or from direct acceptor excitation in the delayed window.

Using these photon-resolved datasets, fluorescence lifetime and anisotropy images can be constructed (**Fig. 1D, left**), and the time-resolved fluorescence intensity (**Fig. 1D, middle**) and anisotropy (**Fig. 1D, right**) can be obtained by summation of the respective photons. The shapes of these time-resolved curves already provide qualitative indications of energy transfer and thus protein interactions.

Quantitative analysis of pixel-wise fluorescence lifetimes, anisotropies, and count rates identifies cellular heterogeneities (**Fig. 1E, left**) and supports the development of biophysical models, including the determination of dimerization association constants (**Fig. 1E, middle**) and the inference of potential interaction sites within MC4R complexes (**Fig. 1E, right**).

### 3.2 Identifying GPCR oligomerization

To identify the self-association potential of our two proteins of interest, we systematically varied the donor to acceptor ratios in titration experiments and performed live-cell FLIM measurements at the basal membrane of HEK293T cells. During transfection, we increased the number of mCherry-tagged receptors while maintaining a constant plasmid DNA quantity. Given that dimer and oligomer formation occur stochastically, this strategy increases the likelihood of forming FRET-active eGFP–mCherry receptor complexes.

In the first analysis step, the fluorescence lifetime images generated by the microscope or exported via open-source software (**Supp. Fig. 1**) can be inspected visually. In this study, we used the open-source software *ttttrlib* (Peulen, et al. 2025) and *ChiSurf* (Peulen 2025) for quantitative analysis. All scripts used to produce the presented results are provided as supplementary material (Greife, et al. 2025). While large lifetime changes are readily detected in fluorescence lifetime images (**Fig. 2, Supp. Fig. 1**), subtle differences or quantitative interpretation require more advanced analysis. Therefore, we defined regions of interest (ROIs) for each cell in each image. We exported the photon arrival times of the ROIs from the green channels, reconstructed time-resolved fluorescence intensity decays, and fitted them with a multi-exponential decay model to extract average fluorescence lifetimes (**eq. 1**; **Fig. 3A**). For both MC4R-A and MC4R-B2, the average eGFP lifetime decreased from approximately 2.4 ns to 1.7 ns as the fraction of mCherry-tagged receptor increased, indicating FRET and thus receptor-receptor interaction.

**Figure 2.**
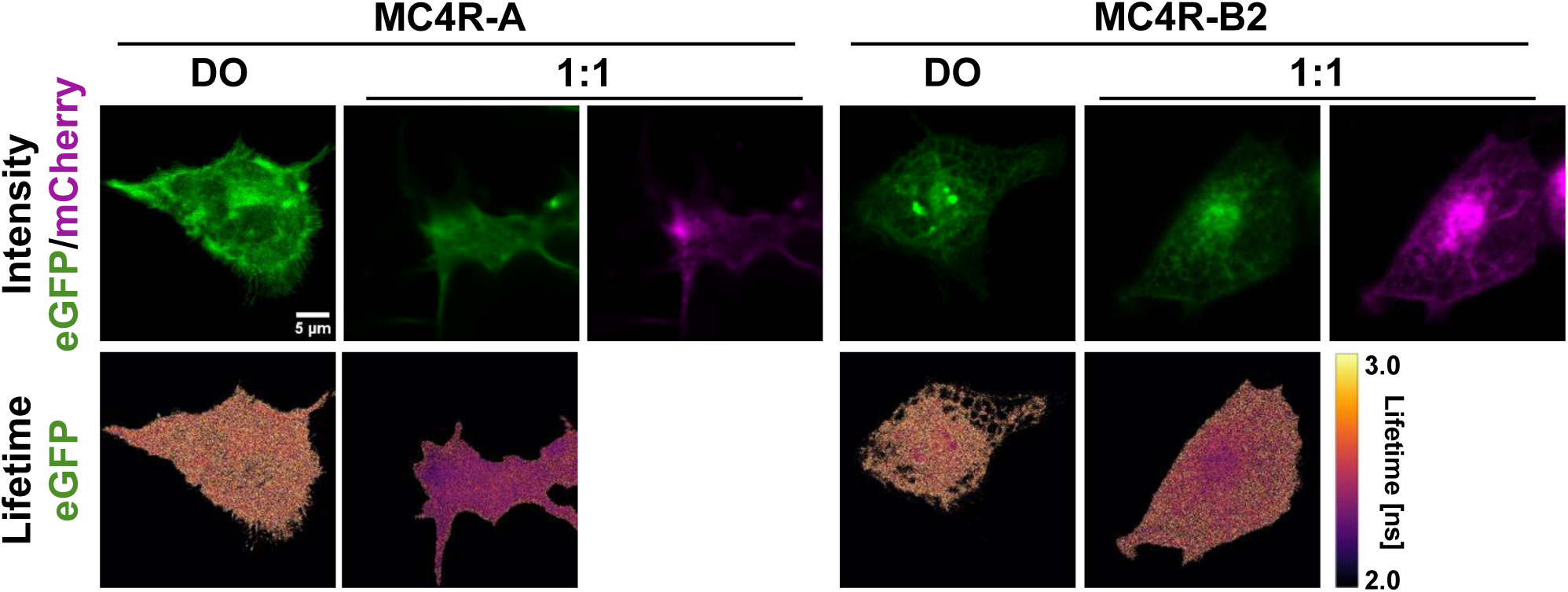
HEK293T cells co-transfected with MC4R eGFP and mCherry-tagged constructs show a decreased average fluorescence lifetime compared to singly transfected cells. (<u>top</u>) eGFP fluorescence intensity in the “prompt” time window of a singly transfected (“DO”) and mCherry fluorescence intensity in the “delay” time window (i.e., direct acceptor excitation) of a co-transfected cell with a 1:1 ratio of eGFP and mCherry plasmid (MC4R-A, left; MC4R-B2, right). (<u>bottom</u>) eGFP fluorescence lifetime image. The total amount of plasmid was kept constant. The average lifetime was computed for pixels with at least 15 photons.

**Figure 3.**
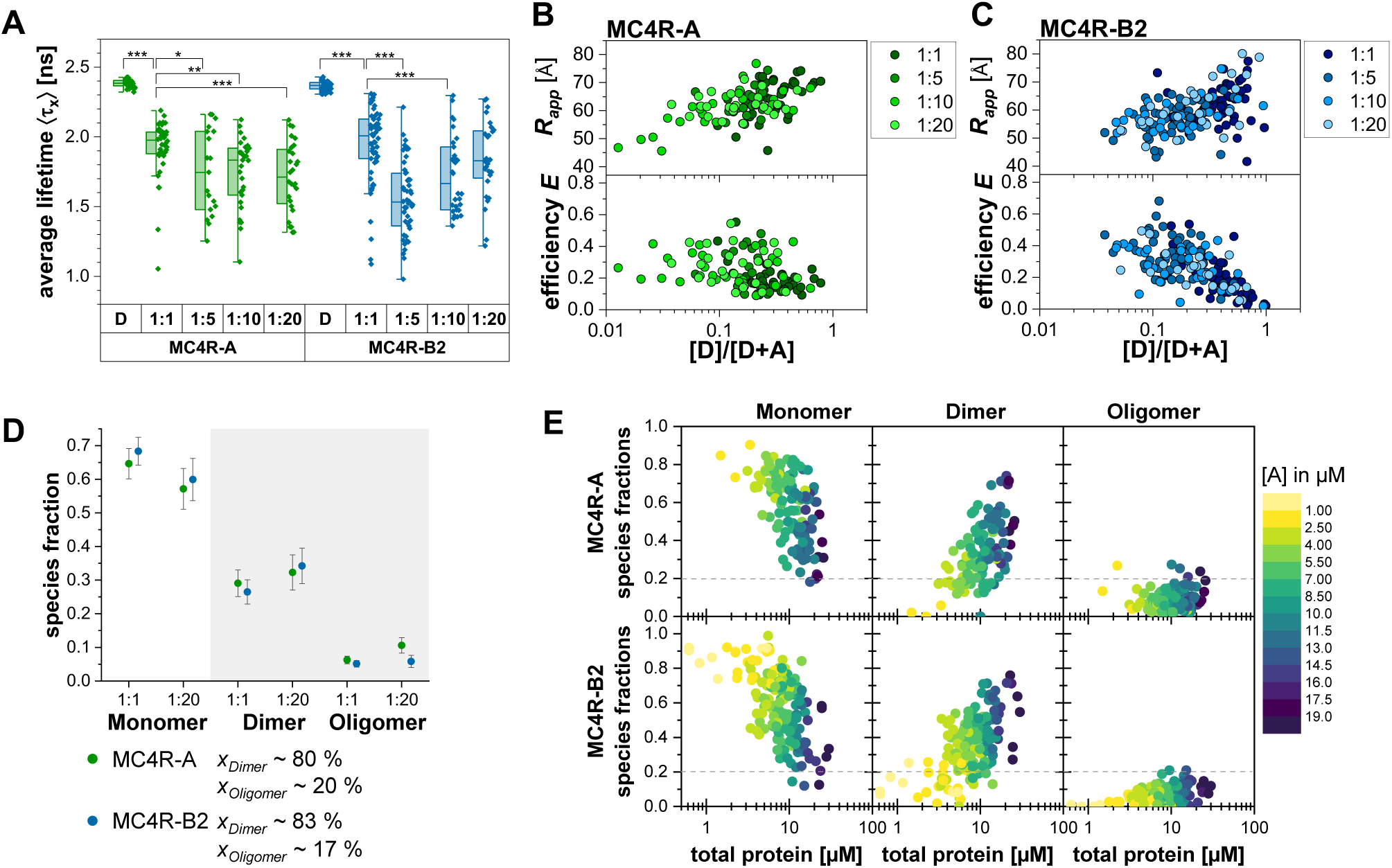
MC4R-A and MC4R-B2 variants show protein concentration-dependent oligomerization. Titration series of eGFP (D) and mCherry (A) tagged MC4R-A (green) and MC4R-B2 (blue) with different D:A ratios. **(A)** The species-weighted average fluorescence lifetime 〈*τ*)_x_ of D at different D:A ratios decreased from ∼2.4 ns to ∼ 1.7 ns with increasing amounts of the acceptor, [A]. **(B-C)** The average apparent inter-fluorophore distance *R_app_* (obtained from the single Gaussian distance fit) and FRET efficiency, *E*, depend on the stoichiometry, [D]/[D+A]. **(D)** Mean and 95% confidence interval of the species fractions from the free two Gaussian distances fit for both MC4R variants based on the DA 1:1 and DA 1:20 data (MC4R-A n=56, MC4R-B2 n=85). **(E)** Global analysis with two Gaussian distances reveals that the species fractions of both MC4R variants depends on the total protein and especially the acceptor concentration, [A] (MC4R-A: n=116; MC4R-B2: n=163; at least four independent experiments). MC4R-mCherry concentrations are color-coded: 0.5 µM (yellow) - 20 µM (dark blue).

Once protein-protein interactions have been confirmed, the next steps depend on the desired level of quantitative detail. The acquired data is higher dimensional. Affinity constants, *K_D_*, can be obtained in multiple ways. For instance, the observed fluorescence lifetimes can be plotted against either the receptor concentration (if available) or a suitable approximation, such as total count rate or the fraction of photons in the prompt window relative to all prompt and delayed photons. This ratio, known as stoichiometry, serves as a proxy for the donor/acceptor composition. In a concentration-dependent dimerization model, the observed FRET efficiency depends on both the total concentration and the acceptor-to-donor ratio. As a result, the slope of FRET efficiency plotted against either parameter already provides initial insights into relative *K_D_* values for the two MC4R variants (**Fig. 3B/C, bottom**).

### 3.3 Quantifying GPCR oligomerization

An accurate and precise quantification of the oligomerization depends on the models’ and reference samples’ accuracy, and the data noise. It is reasonable to assume that the FPs coupled to MC4R adopt multiple conformations. Hence, to accurately interpret our experiments, we analyzed the fluorescence intensity decays using an inter-fluorophore distance model (**eq. 3-5**). The model describes interacting eGFP-mCherry pairs by a Gaussian distribution of width *σ_app_* limited to a physical reasonable range (5-20 Å) and an average distance *R_app_*. The model includes a fraction *x*_*noFRET*_ representing molecules that do not exhibit detectable FRET, comprising both truly monomeric receptors and associated receptors with unfavorable dipole orientations. Molecules not undergoing FRET are either (*i*) non-interacting, monomeric molecules (*x*_DO_) or (*ii*) fluorophore pairs with unfavorably oriented dipoles (*x*_Dipl_). Experimentally, we found that the donor fluorescence lifetime signatures (average lifetime > 2.30 ns) of cells transfected for FRET with very low acceptor expression match the signature of the reference (eGFP-only constructs) cells (average lifetime 2.38 ns (MC4R-A) and 2.36 ns (MC4R-B2), **Fig. 3A**) supporting the accuracy of our reference for analysis. To increase the precision and minimize the data noise, we stabilized our analysis by joint fits across multiple samples. The distribution width *σ_app_* depends on the linker length, linker flexibility, and the local environment (e.g. membrane context).

The extracted distances and the FRET-inactive fractions were plotted against the concentration ratio (**Fig. 3B-C, Supp. Fig. 2A**). For concentration-dependent trend analysis, we approximated *x*_*noFRET*_ as the monomeric fraction, while acknowledging that this represents an upper bound due to orientation-induced FRET silencing. Thus, we neglected *x*_Dipl_ and treated *x*_noFRET_ as fraction of monomeric MC4R, *x_mo_*. When only dimerization occurs, the inter-fluorophore distance remains constant across the concentration range, while *x_mo_* decreases (**Supp. Fig. 2A**). When higher-order oligomers form, the inter-fluorophore distances decrease while *x_FRET_* increases concentration-dependent (Greife, et al. 2016). For MC4R-A, the average inter-fluorophore distance decreased from approximately 70 Å to 55 Å. For MC4R-B2 it decreases from approximately 75 Å to 50 Å (**Fig. 3B–C**). These changes are a clear hallmark for the formation of higher-order oligomers.

These results prompted us to analyze the data with a three-state model comprising monomers, dimers, and oligomers. The dimer contribution was modeled with a Gaussian distance distribution. In the oligomeric state, varying receptor compositions with multiple donor-to-acceptor stoichiometries form. An eGFP-tagged receptor may be surrounded by one or more mCherry-tagged receptors, with each acceptor contributing independently to donor quenching. The collective FRET from multiple acceptors produces a notably high apparent FRET-rate constant, *k_FRET_* (Bunt and Wouters 2017). This can be represented as an effectively short donor–acceptor distance, even though the actual distances in an oligomer may be longer depending on the complex architecture. For computational tractability, we approximated the oligomeric *k_FRET_* with a short distance and a narrow distribution (*σ_ol_* = 1 Å).

Fitting was performed in two rounds: first the dimer, *R_di_*, and oligomer distance, *R_ol_*, were allowed to vary freely (**Fig. 3D, Supp. Fig. 2B-C**), and then with distances fixed to the average values obtained in round 1 to stabilize the fits (**Supp. Fig. 2D**). For MC4R-A we obtained an average *R_di_* of 60.0 ± 7.2 Å (mean ± SD) and *R_ol_* of 37.4 ± 3.2 Å, and for MC4R-B2 we obtained an average *R_di_* of 60.1 ± 7.5 Å and *R_ol_* of 37.3 ± 4.2 Å, i.e. both MC4R-A and MC4R-B2 show identical inter-fluorophore distances. Finally, we plotted the resulting species fractions of the fixed distances model - monomer, dimer, and oligomer - as a function of total protein concentration, color-coded by acceptor expression (**Fig. 3E; Supp. Fig. 2E-J**). As expected, the monomer fraction decreased with increasing concentration, whereas the dimer fraction increased. The oligomer fraction increased modestly but consistently, reaching a maximum of 20 % for both variants.

Taken together, these results indicate that, in addition to robust dimerization, both MC4R-A and MC4R-B2 form a minor population of higher-order oligomers in live cells.”

### 3.4 Segmentation extends concentration ranges in live-cell experiments

Using a titration series with varying D to A ratios, 1:1 to 1:20, we probed receptor concentrations up to ∼50 µM, with acceptor concentrations reaching ∼22 µM. However, at donor-to-acceptor ratios of 1:10 or higher, it became increasingly difficult to identify cells that expressed both donor- and acceptor-tagged constructs. Notably, proteins in living cells are typically not evenly distributed; instead, they accumulate locally in specific organelles or, in the case of GPCRs, concentrate in endocytic or recycling vesicles (Eichel and von Zastrow 2018) or membrane nanodomains (Calebiro and Grimes 2020, Calebiro et al. 2021). These heterogeneous concentration patterns can be leveraged into an analytical advantage, as they naturally extend the accessible concentration range and reduce intracellular averaging, thereby enhancing the discriminative power for quantifying monomers, dimers, and higher-order oligomers.

Here, we present two approaches for subdividing a single-cell membrane ROI into distinct subregions (**Fig. 4A; Supp. Fig. 3**). The first approach is intensity-based, while the second relies on the Number&Brightness (N&B) method (Digman, et al. 2008) (**Supp. Fig. 4**), which evaluates pixel-wise fluorescence fluctuations across the acquired image series (50 frames per measurement). Segmenting the images extends the concentration range and reduces the number of measurements.

**Figure 4.**
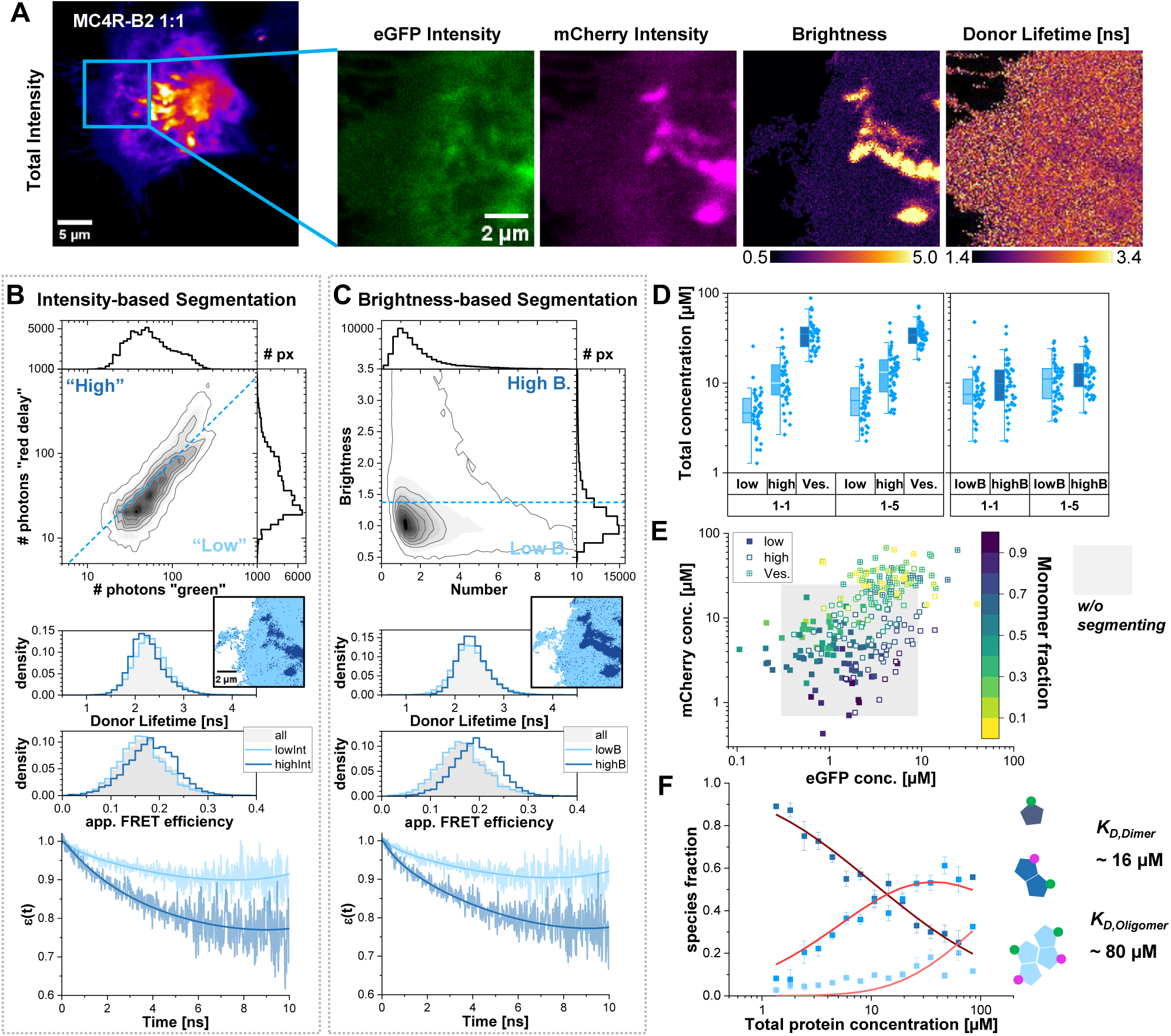
Intensity and brightness-based cell segmentation expands the accessible concentration range by intra-cell heterogeneities illustrated for MC4R-B2. **(A)** Donor fluorescence lifetimes, brightness and intensity images in HEK293T basal plasma-membranes co-transfected with MC4R-B2-eGFP and mCherry highlight heterogeneities (green: eGFP fluorescence intensity, magenta: directly excited mCherry intensity) **(B)** Intensity-based segmentation: 2D pixel-histograms of the green and red-delay photon counts with marginal distributions. Pixels above the dashed line were assigned as “high” and pixels below the line as “low” intensity region. The middle panel shows the pixel-wise mean donor fluorescence lifetime and intensity-based apparent FRET efficiency (light grey) for the whole cell, the low-intensity regions (light blue), and high-intensity regions (dark blue). The bottom panel shows the corresponding FRET-induced donor decay. **(C)** Same as (B) for the brightness-based segmentation with a threshold level of *B* = 1.375. **(D)** Total concentrations were obtained after segmenting each cell into three intensity regions (low, high, vesicle) or the brightness regions (lowB, highB). **(E)** Donor vs acceptor concentration for the intensity-based sub-segmentation (low: filled symbol; high: open symbol, vesicle: crossed symbol), color-coded by the fraction of monomers (from global analysis). The grey-shaded area indicates the concentration range probed without the sub-segmentation. **(F)** Average monomer (dark blue), dimer (blue), and oligomer (light blue) fractions *vs* total protein concentrations. The protein concentrations were grouped into logarithmically spaced bins. The corresponding species fractions were fitted with the oligomerization model (dark red, red. and light red, **eq. 17-19**). The error bars are the standard error of the means. MC4R-A results are shown in the **Supp. Fig. 5**.

In our intensity-based segmentation strategy, demonstrated for a MC4R-B2 cell, we construct pixel-wise 2D histograms of green/donor photon counts versus red photons detected after direct acceptor excitation (“red delay”) (**Fig. 4B, top**) that are divided into “low” and “high” regions. The “high” region emits more acceptor photons compared to the “low” region. As expected, the “high” region displays shorter donor fluorescence lifetimes, *τ*, and higher intensity-based apparent FRET efficiencies, *E_app_*, in pixel-wise distributions (**Fig. 4B, middle**). Likewise, the increase of FRET in “high” regions is highlighted by the time-dependent depopulation of the donor’s excited state in the FRET-induced donor decay *ε*(t) (Peulen, et al. 2017) (**Fig 4B, bottom**). In an *ε*(t) analysis, the donor decay in the presence of an acceptor, *f_DA_(t)*, is divided by the reference donor-only decay, *f_DO_(t)*. The slope of *ε*(t) reflects the FRET-rate constant, while its offset corresponds to the fraction of non-interacting molecules *x_noFRET_* (monomer/unfavorable oriented dimer dipoles). In the “high” region, *x_noFRET_* decreased to ∼0.75 compared to ∼0.9 in the “low” region, confirming the increased proportion of interacting receptors. In our fully automated intensity-based analysis workflow, we first discriminated rapidly moving vesicles, which contain densely packed receptors, and treated them as separate pixel class (**Supp. Fig. 3**). The remaining membrane region was segmented automatically into “high” and “low” intensity subregions using Otsu thresholding; and donor fluorescence intensity decays were extracted for each.

In the second N&B-based segmentation strategy, we compute the intensity mean and variance over pixels. The apparent brightness *B* is the ratio of variance to mean intensity. The apparent number *N* is the total intensity divided by *B* (Digman, et al. 2008). Monomeric fluorophores exhibit *B* ≈ 1 (**Supp. Fig. 4**). A 2D *N* vs. *B* histogram (**Fig. 3C, top**) reveals distinct populations corresponding to different molecular brightness. Pixels with *B* > 1.375 were classified as “highB”, while all others were assigned to “lowB”. As expected, “highB” pixels were enriched in regions containing a larger fraction of dimeric or oligomeric receptors, consistent with the trends observed in the intensity-based segmentation (**Fig. 4C, middle/bottom**).

Fluorescence intensities and donor decays were exported for the five different pixel categories - “low” intensity, “high” intensity, “vesicles” (very high intensity), “lowB”, and “highB” (brightness) - from D:A samples transfected at 1:1 and 1:5 ratios. We calculated the corresponding protein concentrations (**Fig. 4D, Supp. Fig. 5A**). As anticipated, “high” and “vesicles” regions contained elevated protein levels relative to “low”, whereas “lowB” and “highB” showed only small differences in concentration. Depending on the biological question, vesicles may be of particular interest - for example, in studies of GPCR activation or trafficking - or may be excluded. In the subsequent analysis, we included vesicles as high-concentration regions. All subregions were analyzed using the previously established three-state model (monomer-dimer-oligomer) with fixed dimer and apparent oligomer donor-acceptor distances. Donor versus acceptor concentrations were plotted and color-coded by monomer fraction (**Fig. 4E**). For the intensity-based segmentation of MC4R-B2, a clear decrease in monomer fraction was observed in high concentration regions, accompanied by increased dimer and oligomer fractions (**Supp. Fig. 5B-F**). Importantly, the intensity-based segmentation extended the accessible concentration range and ratios - indicated by the grey rectangle in **Fig. 4E** and **Supp. Fig. 5** - thus facilitating *K_D_* estimation without requiring extreme transfection ratios (and reducing the number of required live-cell experiments).

To determine affinity constants, the species fractions were grouped by total protein concentration into 15 logarithmically spaced bins and fitted using an oligomerization model (**Fig. 4F**, **eq. 17-19**) to yield *K_Dimer_* of ∼16 µM for MC4R-B2 and ∼20 µM for MC4R-A (**Supp. Fig. 5G, Supp. Table 2**). The *K_Oligomer_* for both variants exceeded the measured concentration range (>80 µM). Note that fitting the non-binned (**Supp. Fig. 5H-I**) or the full ROI data (**Supp. Fig. 6**) results in similar values (**Supp. Table 3**). In contrast, while the N&B-based segmentation successfully identified pixels with elevated apparent brightness (“highB”), the resulting expansion of accessible protein concentration range was relatively small compared to the intensity-based segmentation (**Supp. Fig. 7**). Consequently, in this particular example the N&B approach provided only a limited additional leverage for estimating oligomerization equilibria. The intensity-based segmentation produced a substantially wider distribution of protein concentrations, which in turn improved the robustness of the *K_D_* estimation. Nonetheless, N&B may be more powerful in systems with larger brightness heterogeneity more strongly reflecting underlying oligomeric states.

### 3.5 Identifying interactions with single fluorescent protein tags

So far, we focused on FRET between spectrally distinct fluorophores (heteroFRET). Energy transfer between identical fluorophores - a process known as homoFRET - can be used in combination with heteroFRET to determine whether protein A interacts exclusively with itself (homoFRET) or additionally with protein B (heteroFRET). In contrast to heteroFRET, homoFRET does not affect the fluorescence lifetime of the donor; instead, it manifests as a reduction in fluorescence anisotropy (Liput et al. 2020, Vogel, et al. 2015, Vogel, et al. 2009). Fluorescence anisotropy compares the difference between parallel and perpendicular emission intensities following excitation with linearly polarized light. The resulting steady-state anisotropy, *r_ss_*, (based on fluorescence intensities, **eq. 11**) and time-resolved anisotropy, *r*(*t*), (based on the time-resolved fluorescence decays) reflects the rotational mobility of the fluorophore. Rapidly tumbling molecules exhibit greater depolarization and thus lower anisotropy values. Fluorescent proteins in solution typically have steady-state anisotropies around ∼0.33, while fluorescent proteins fused to membrane proteins - such as GPCRs - exhibit higher values due to restricted rotational diffusion.

Like the heteroFRET analysis, the homoFRET workflow proceeds stepwise: (*i*) Determine the apparent (uncorrected) steady-state anisotropy per cell and plot it against concentration or count rate, (*ii*) determine polarization-dependent correction factors for the instrument, (*iii*) examine the time-resolved fluorescence anisotropy decays to identify fast components, and (*iv*) fit the time-resolved anisotropy decays using appropriate rotational or energy-transfer models.

Steady-state anisotropy images (**Supp. Fig. 8**) visually highlight qualitative differences between dim and bright cells (expression level). Steady-state anisotropies depend on the tumbling of the fluorophores that can be affected by their local environment. Freely and bound rotating FPs with a lifetime of 2.4 ns and rotation times of 16 ns and 100 ns have steady-state anisotropies of 0.33 and 0.37, respectively. Notably, we find a higher anisotropy for membrane anchored FPs in MC4R-A compared to non-membrane anchored FPs in MC4R-B2 (**Fig. 1A**).

A quantitative analysis of absolute anisotropies requires careful attention on the instrumental calibrations. Upon polarized excitation, homoFRET introduces an additional depolarization pathway, reducing the observed anisotropy. For accurate analysis, the additional depolarization in high-NA objectives must be quantified and corrected (**Supp. Fig. 9; Methods**) (Erdelyi, et al. 2014, Koshioka, et al. 1995, Liput, et al. 2020). We quantified the steady-state anisotropy *r*_ss_ (**eq. 11**) for both DO samples and for the directly excited acceptor signal (“red delay”) in the D:A-transfected cells (1:1 and 1:5; **Fig. 5A,B**). Please note that the concentration refers to the total protein concentration (D+A) as the monomer-dimer-oligomer equilibrium is also influenced by the (invisible) donor-tagged receptors in the same cell (For reference, **Supp. Fig. 10A,B** show the same data using only the acceptor concentration). At comparable concentrations, MC4R-B2-eGFP exhibited lower *r*_ss_ values and weaker concentration dependence than MC4R-A-eGFP. For mCherry, the anisotropy depended strongly on acceptor concentration, again with slightly lower values for MC4R-B2 compared to MC4R-A at matched expression levels (**Fig. 5A-B**).

**Figure 5.**
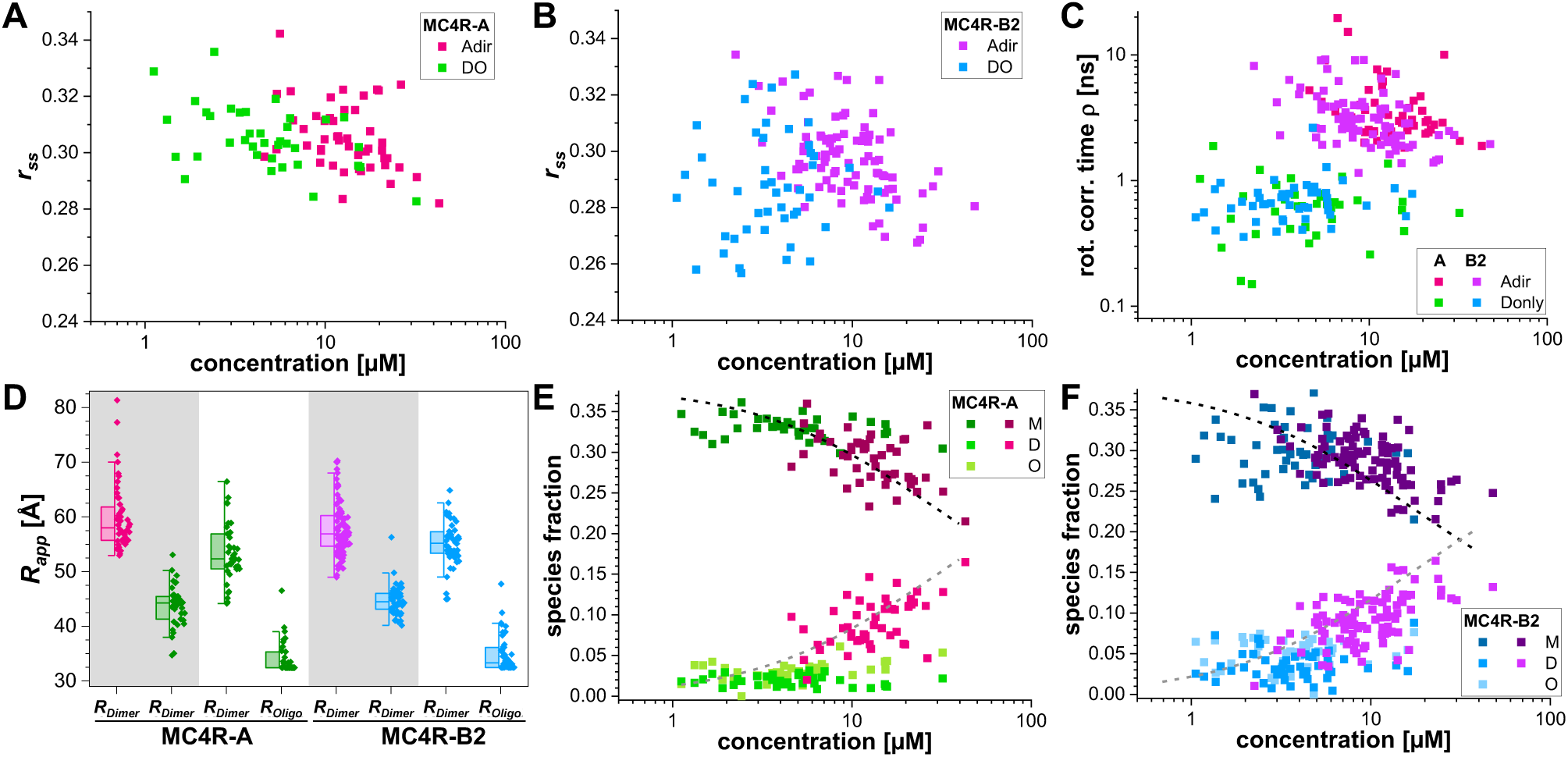
Single-color experiments: HomoFRET detects dimerization and oligomerization. **(A)** Steady-state anisotropy of eGFP in the absence of acceptors (DO, green) and directly excited acceptor mCherry (direct Acc., magenta) in MC4R-A transfected cells. Concentrations are the total protein concentrations**. (B)** Corresponding MC4R-B2 experiments (DO cells are blue; co-transfected cells are violet). **(C)** Anisotropy relaxation times of the dimer model (**eq. 6**) of the cells shown in (A, B). **(D)** Apparent inter-fluorophore distance, *R_app_*, computed for the relaxation times of MC4R-A and MC4R-B2 (grey background, dimer model; magenta/violet, direct acceptor excitation; green/blue, DO). DO samples were described by the oligomer model (**eq. 7**; white background). **(E)** Species fractions (global, fixed FRET-rate constants, oligomer model) of the MC4R-A DO samples (green shades) and of the directly excited acceptors (magenta shades, dimer model, global, fixed FRET-rate constants) (M= monomer, D=Dimer, O=Oligomer). For DO, oligomer and dimer fractions were combined. The combined acceptor and DO data are described by a dimerization binding isotherm (black: Monomer, gray: Dimer). (**F)** Same as (E) for the MC4R-B2 variant.

To jointly interpret heteroFRET and homoFRET of eGFP and mCherry data, differences in the Förster radius, *R*_0_, must be considered, which define the accessible distance range. For eGFP-mCherry, eGFP-eGFP, and mCherry-mCherry *R*_0_are 52 Å, 50 Å, and 44 Å, respectively (Lambert 2019). In the FRET-sensitive distance range of roughly 0.5–1.5 *R*_0_ (Hellenkamp et al. 2018), we probe distances up to ∼67 Å for an mCherry-mCherry and up to ∼78 Å for an eGFP-mCherry pair. The effective FRET-rate constant is doubled in homoFRET due to the bidirectional energy transfer (Liput, et al. 2020). Consequently, in oligomeric states, lifetimes can fall below the instrument response function width, typically 200–500 ps. A qualitative indicator for fast depolarization is a drop of the initial anisotropy decay amplitude, *r*_0_, using the anisotropy decay of an FP control (*r*_0_≈ 0.38) as a reference. Fast depolarization processes such as homoFRET reduce *r*_0_ (**Supp. Fig. 9F)** and reflect in steady-state anisotropy images and time-resolved anisotropy decays for three MC4R-A-eGFP cells (**Supp. Fig. 8A**). For MC4R-A, we find a correlation between a steady-state anisotropy reduction and a drop in *r*_0_ compared to the eGFP references. We observed a similar trend for MC4R-B2-mCherry in co-transfected eGFP and mCherry cells (**Supp. Fig. 8B**). Both are a clear qualitative indicator for MC4R oligomerization.

### 3.6 Quantifying interactions with single fluorescent protein tags

We process the homoFRET data independently of the heteroFRET data, not using knowledge of the oligomerization to compare the outcomes of the two methods. We analyze time-resolved anisotropy homoFRET data using a dimer and an oligomer model, as *a priori*, it is unclear whether a biomolecule dimerizes or forms oligomers. Analogous to the heteroFRET approach, we performed the analysis in two rounds. In the first round, samples were analyzed individually to obtain an average homoFRET-rate constant (**Supp. Fig. 10C,D**). In the second round, to stabilize the analysis, species amplitudes were the sole variable parameters. Our monomer model has one relaxation time for the global rotation, while dimer and oligomer mixture models have 2 and 3 relaxation times. Membrane proteins rotate slowly compared to the fluorescence lifetime (Balakrishnan et al. 2022). Thus, the rotational component was fixed to 100 ns.

When analyzing the data using dimer models in the first round (individual analysis of samples), we observed an increase of the fast relaxation amplitude and a drop in the relaxation time from ∼10 ns to ∼1 ns (**Fig. 5C**) along with increasing acceptor concentrations. Surprisingly, in DO samples, the fast relaxation time remained nearly constant (∼0.8 ns) (**Fig. 5C**). The concentration-independent behavior for eGFP was unexpected. A close inspection of the model parameters revealed that the perpendicular channel contained substantial short-lived “scatter” components (**Supp. Fig. 10E,G**). For mCherry the scatter contribution was considerably lower and showed a linear relationship between parallel and perpendicular channels (**Supp. Fig. 10F,H**). These observations suggested a missing anisotropy amplitude (as seen in **Supp. Fig. 8-9**) was compensated/masked by “scatter”. Therefore, we reanalyzed both eGFP and mCherry data of DO and co-transfected datasets, respectively, by our oligomer model (**Supp. Fig. 10I-L**). In the oligomer model, oligomers result in a drop of the time-resolved fluorescence anisotropy at short times. Thus, instead of being described by scattered light, eGFP data is characterized by oligomer formation **(Supp. Fig. 10E,G**). For mCherry, contrary to eGFP, we found low oligomer fractions (**Supp. Fig. 10J,L**), likely due to the shorter *R*_0_ of mCherry.

To compare heteroFRET and homoFRET, the obtained relaxation times were converted to FRET-rate constants and apparent distances (**eq. 6-10, Fig. 5D**). In the dimer model, we obtained average distances of ∼60 Å for MC4R-A-mCherry (*ρ* ∼4.32 ns), ∼44 Å for MC4R-A-eGFP (*ρ* ∼0.74 ns), ∼57 Å for MC4R-B2-mCherry (*ρ* ∼3.33 ns), and ∼45 Å for MC4R-B2-eGFP (*ρ* ∼0.72 ns). The oligomer model for MC4R-A-eGFP yielded distances of 53 Å (*ρ*_*di*_ ∼3.09 ns, dimer) and 34 Å (*ρ*_*ol*_ ∼ 0.22 ns, oligomer), while MC4R-B2 produced 55 Å (*ρ*_*di*_ ∼2.58 ns) and 35 Å (*ρ*_*ol*_ ∼0.19 ns), respectively. For the oligomer model, the apparent distances match the heteroFRET-derived distances, whereas the dimer distances appeared shorter compared to the heteroFRET distances. To further compare homo- and heteroFRET, we determined species fractions to estimate homoFRET affinity constants (**Fig. 5E-F**). The pooled eGFP and mCherry data show a clear decrease of monomers with a concomitant increase in dimers/oligomers. A dimer model fitted to the pooled data of MC4R-A and MC4R-B2 estimates *K_Dimer,app_* of 56 µM and 31 µM, respectively (**Supp. Table 4**). Consistent with the heteroFRET data, we find a higher dimerization affinity for MC4R-B2. This demonstrates how homoFRET can complement heteroFRET experiments.

### 3.7 Homo- and HeteroFRET occur in parallel in the co-transfected samples

We processed homo- and heteroFRET data independently; however, as dimers and higher-order oligomers form stochastically, both homo- and heteroFRET occur in parallel. HomoFRET affects the fluorescence anisotropy while heteroFRET primarily shortens the donor lifetime (**Fig. 6A**). To visualize the simultaneous contribution of homo- and heteroFRET in our samples, we plotted steady-state anisotropies against the donor fluorescence lifetime for all DA and DO cells. As expected, the formation of dimers and higher-order oligomers led to concurrent changes in fluorescence lifetime and anisotropy: heteroFRET shortens the donor lifetime, homoFRET reduces the observed anisotropy.

**Figure 6.**
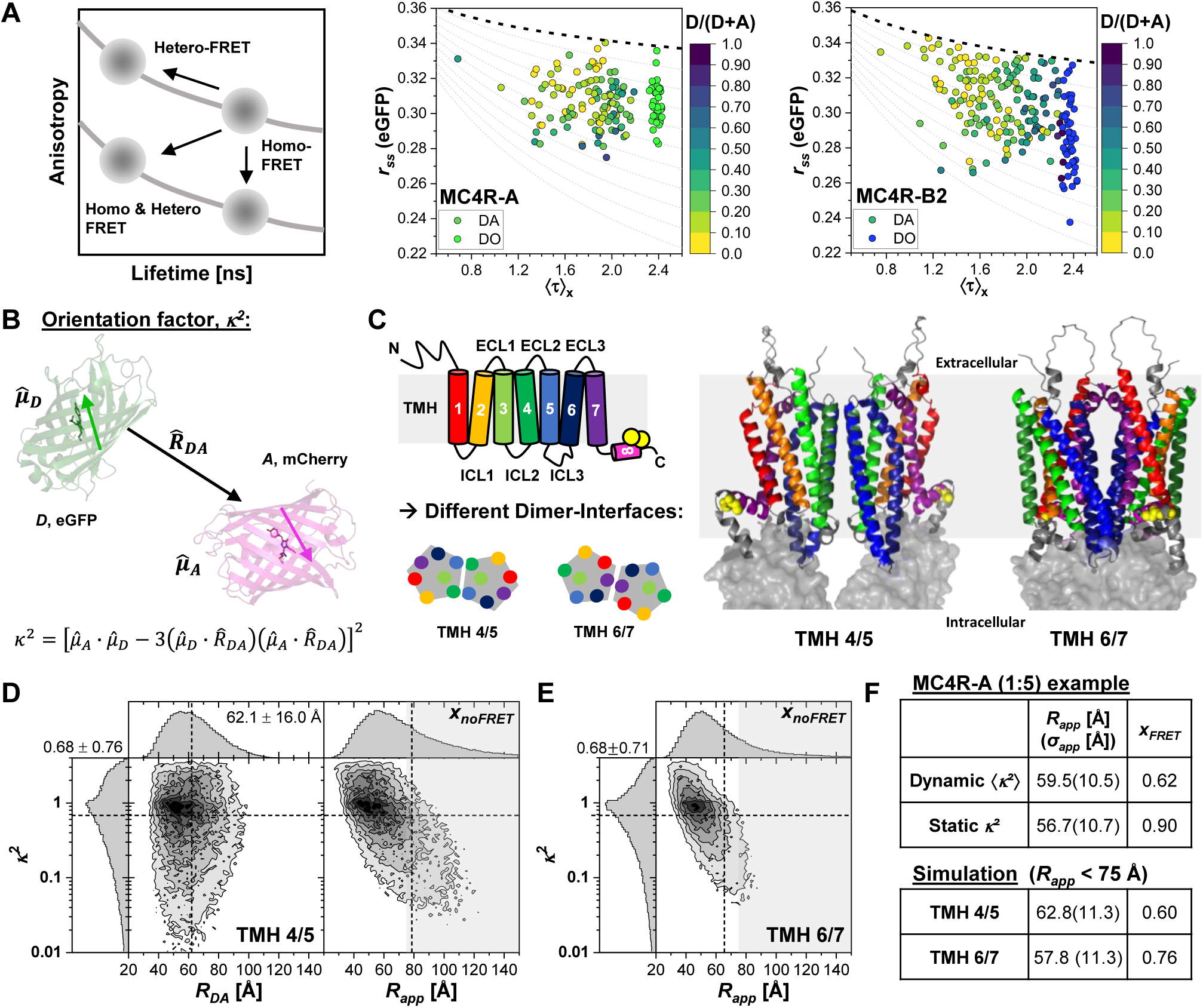
HomoFRET and heteroFRET occur simultaneously and simulation-guided analysis can help defining the interaction interface. **(A)** Average fluorescence lifetime and observed steady-state anisotropies are related by the Perrin-equation (grey, *r*_*obs*_ /*r*_0_ = 1 + *τρ*^−1^). HeteroFRET reduces the lifetime and increases the anisotropy. HomoFRET reduces anisotropy without altering the fluorescence lifetime (left). Scatter plot combining average lifetimes and steady-state anisotropies of MC4R-A (middle) and MC4R-B2 (right). Data points from co-transfected experiments are color-coded by stoichiometry, shown as [D]/[D+A]. DO samples are shown in green (MC4R-A) and blue (MC4R-B2). Black and grey lines show the bi-exponential Perrin-equation with 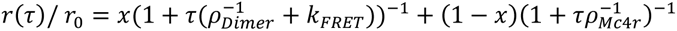 where *x* is the fraction of a fast component (increasing from 20 – 65 % in 5 % steps), and *k*_*FRET*_ = 0.62 ns^−1^ (average FRET-rate of the dimer model). **(B)** The orientation factor *κ*² depends on the mutual donor and acceptor dipole orientation defined by the unit vectors *μ̂*_*D*_, *μ̂*_*A*,_ and *R̂*_*DA*_. (**C**) AlphaFold3 (Abramson, et al. 2024, Jumper, et al. 2021) predicts different dimerization interfaces for MC4R-A. ICL: Intracellular loop; ECL: Extracellular loop; TMH: Trans-membrane helices; N: N-terminus, C: C-terminus; yellow circles: membrane anchor. Dimerization via TMH 4/5 (left) or TMH 6/7 (right). eGFP and mCherry are shown as semitransparent surfaces, the grey-shaded area indicates the cell membrane. **(D)** Simulated inter-fluorophore distances, *R_DA_*, orientation factors *κ*², and resulting apparent distances *R_app_* of FPs attached to MC4R-A for the TMH4/5 dimer. Dashed lines: Mean values for *κ², R_DA,_* and *R_app_*, numbers in the 2D histogram report the mean and width of the distributions. (**E**) Simulated apparent inter-fluorophore distances for the TMH6/7 dimerization model. **(F)** MC4R-A cell data (1:5 transfection) fitted by the Gaussian distance distribution model (**eq. 3-5**) for a single average *κ*² = 2/3 and an isotropic *κ*² distribution; bottom: distribution mean and width of TMH4/5 and TMH6/7 dimer simulations by assigning all models with *R_app_* > 75 Å to *x_noFRET_*. A detailed simulation workflow and results for the TMH3/4 and TMH5/6 model are shown in **Supp.** Fig. 11.

When visualizing experiments in a two-dimensional anisotropy-lifetime space, as expected, only a set of Perrin curves 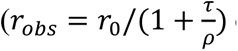 can describe the data, as the downward shift driven by homoFRET and an upward/leftward shift driven by heteroFRET-induced lifetime reductions happen simultaneously to a different extent (**Fig. 6A**). However, within the data clear systematic trends are observed for both MC4R-A and MC4R-B2, highlighted by a color-coding of the stoichiometry shown as the ratio of donor to total protein concentration (**Fig. 6A, middle/right**). With decreasing donor ratio, the DA samples shift towards shorter lifetimes and lower anisotropy values, a hallmark for homo-and heteroFRET. To guide the visualization, we have added a set of Perrin lines for slow rotating molecules with varying degrees of homoFRET. With decreasing donor fraction, the data points shift to Perrin lines with a higher fraction of fast depolarization (top black line: 20 %, bottom grey line: 65 %), confirming that both processes occur in parallel in the cellular environment.

Although in the present experiments both eGFP and mCherry were fused to the same receptor variant, the observed spread of points in the anisotropy-lifetime space illustrates the sensitivity of this approach to mixed interaction modes: In co-expression experiments of distinct receptor variants (e.g., MC4R-A-eGFP with MC4R-B2-mCherry), the relative contribution of homotypic (A-A, B2-B2) versus heterotypic (A-B2) interactions could be estimated as a function of the respective donor and acceptor concentrations.

### 3.8 Accounting for orientational κ² effects

The observed FRET efficiency, *E*, is determined by three main factors: (*i*) the inter-fluorophore distance *R*_*DA*_, (*ii*) the spectral overlap between donor emission and acceptor excitation (determining the Förster radius together with donor fluorescence quantum yield), and (*iii*) the relative orientation of the donor and acceptor transition dipole moments, *κ*² (Förster 1948) (**Fig. 6B**). Notably, the influence of *κ*² is often neglected as it is experimentally challenging to determine (Hummer and Szabo 2017, Meer et al. 2013). Two limiting regimes can be distinguished: in the dynamic averaging regime, fluorophores rotate fast compared to their fluorescence lifetime (rotation ≪ τ), allowing their orientations to average (Dale et al. 1979, Peulen, et al. 2017). In the static regime fluorophores rotate slowly compared to the fluorescence lifetime and thus requires considering a *κ*² distribution (Hummer and Szabo 2017, Meer, et al. 2013, Peulen, et al. 2017). For intensity-based FRET, an average ⟨***κ***^2^⟩ of 0.476 should be used (Meer, et al. 2013). FPs attached to a protein can be sterically restrained. Thus, fluorescence analysis benefits from molecular simulations to assess spatial and *κ*² distributions.

As an example, we simulate the *κ²* distribution of a heteroFRET pair by feeding the sequences of MC4R-A-eGFP and MC4R-A-mCherry into Alphafold3 (AF3) (Abramson, et al. 2024) and build the dimer models (Evans et al. 2022). AF3 predicts models with four distinct dimerization interfaces. Dimerization occurred either via transmembrane helix (TMH) 4/5, TMH6/7, TMH5/6 or TMH3/4 (**Fig. 6C**). Notably, for GPCRs, all these different interaction interfaces have been found (Breitwieser 2004). To obtain distributions over inter FP distances and orientation factors we coarse grain the MC4R-FP dimers in the integrative modelling platform (IMP) (Russel, et al. 2012) and sample the sterically allowed conformational space by Markov-Chain-Monte-Carlo (**Supp. Fig. 11**). For the obtained models, we compute inter-fluorophore distances, *R_DA_* and *R_app_*, and orientation factors (**Fig. 6C-E, Supp. Fig. 11**). The approach to coarse grain a sample conformational spaces in IMP was implemented in open-source software, FPSIMP (https://github.com/fluorescence-tools/fpsimp) While in all models the average 〈*κ*²) only slightly deviated from 2/3, the broad *κ*² distribution highlights its influence on *R_app_* (**Fig. 6D-E, Supp. Fig. 11**).

Even though one may expect steric restraints imposed by the linkage and the membrane, we find a good agreement between the peak positions of apparent distance, *R_app_*, as compared to the distance distribution, *R_DA_* **(Fig. 6D**). This agreement supports our analysis by a simple distribution model, which does not explicitly account for the orientation factor distribution. Note, *R_DA_*, is the physical distance, while *R_app_* is an apparent distance that originates from the assumption that 〈*κ*²) = 2/3. The main discerning feature between *R_DA_* and *R_app_* distributions is the tail towards longer apparent distances. A direct fluorescence decay analysis of MC4R-A 1:5 transfected cell by a single Gaussian distribution (**eq. 3-5**) in the dynamic and static regime (**Fig. 6F**) supports this finding: the inter-fluorophore distances change marginally from 60 Å to 57 Å while *x_FRET_* increases from 62 % to 90 %. The difference in the fraction of FRET-active species is explained by the explicit handling of “missing/invisible” fraction of FRET-molecules due to unfavorably oriented dipoles. While fast rotating fluorophores adopt conformations favorable for FRET, slowly rotating fluorophores are “stationary”. Thus, orientational factors can prevent energy transfer, even at short distances. Accordingly, our definition of *x*_noFRET_ throughout the study included both monomers and dimers/oligomers with unfavorable dipole orientations. Consequently, while fitting with *κ*² distributions could slightly shift the estimated *K_D_*, the overall interpretation of inter-fluorophore distances and oligomerization trends remains robust.

### 3.9 Live-cell FRET experiments inform on dimerization interfaces

The main objective in the design of the MC4R variants was to quantify the different dimerization degrees. Motivated by the agreement between the simulations and the experiments, we use the FRET data to study dimerization interfaces. Clearly, only given the FRET data alone, this is an underdetermined problem. However, given the FRET data and different competing models, these models can be ranked by their agreement with the experiments.

For a rigorous comparison with defined statistics, the fluorescence decays and the models are compared directly considering the data shot-noise and other experimental uncertainties (Greife, et al. 2016). In this analysis, we compare only the FRET-active species with *R_app_* < 75 Å and assign all other structural models to *x*_noFRET_. Next, we compute average apparent distances, ⟨*R*_*app*_⟩, and distribution widths. For TMH4/5 we obtain an average of 63 Å with a width of 11 Å. The other three competing models TMH6/7 (**Fig. 6E**), TMH1/7 (including helix 8), and TMH3/4 (**Supp. Fig. S11B,C**) show all very similar distance distributions of the FRET-active fraction with mean and width of 58 ± 11 Å, 57 ± 12 Å, and 59 ± 12 Å, respectively. Notably, point mutations in ICL2 and adjacent TMH3/4 region strongly reduce the dimerization (Piechowski, et al. 2013). Thus, the TMH 4/5 dimer simulations, in which the ICL2 and adjacent cytosolic TMH3 residues form the dimerization interface, agree best with our experiments (**Fig. 6D**). In the TMH3/4 model ICL2 and intracellular residues of TMH3 are far apart. This shows that a combination of biochemical studies, live-cell FRET experiments, and structural simulations can delineate receptor arrangements and inform on dimerization interfaces.

## 4 Discussion

### 4.1 PIE-FLIM as a powerful tool to study protein–protein interactions in live cells

Disentangling the intricate interactions of biomacromolecules in living cells remains challenging and requires methods that provide nanoscale spatial sensitivity and quantitative spectroscopic readouts while remaining compatible with live-cell imaging. In this context, live-cell PIE-FLIM proved to be a particularly powerful approach. FRET combined with FLIM provides a uniquely powerful framework to study nanoscale protein-protein interactions under near-physiological conditions. By reporting on molecular proximities below the diffraction limit, these techniques are ideally suited for investigating dynamic receptor organization, structural plasticity, and state-dependent oligomerization in living cells (Greife, et al. 2016, Kravets, et al. 2016, Weidtkamp-Peters, et al. 2009, Zhang, et al. 2025). For complex assemblies such as GPCR signaling platforms, where ligand-induced rearrangements and environment-driven interactions constantly remodel the nanoscale landscape, PIE-FLIM enables quantitative and time-resolved interrogation of molecular architecture (Greife, et al. 2016, Kravets, et al. 2016, Somssich et al. 2015, Weidtkamp-Peters, et al. 2009). For GPCRs such as MC4R, ligand binding can shift the equilibrium of oligomeric states, rearrange protomer interfaces, and alter both inter- and intramolecular FRET signatures (Latorraca et al. 2017).

### 4.2 Both MC4R-A and MC4R-B2 show modest but significant oligomerization

Here we demonstrated the use of live-cell spectroscopy by re-examining the homotypic protein interactions of MC4R. Previous work has established that MC4R forms dimers (Kleinau, et al. 2020, Liu, et al. 2023, Nickolls and Maki 2006) and proposed possible interaction interfaces (Piechowski, et al. 2013), our data extends this findings by demonstrating that MC4R populates a broader spectrum of oligomeric states, ranging from monomers to higher-order assemblies. This was suggested previously (Chapman and Findlay 2013), and we now provide quantitative estimates of the corresponding affinity constants, moving beyond the largely qualitative or binary interaction assessments reported so far.

Both MC4R-A and MC4R-B2 displayed modest but reproducible oligomerization, with MC4R-B2 consistently exhibiting a slightly higher apparent affinity than MC4R-A. This trend was independently supported by both homoFRET and heteroFRET-based analysis, lending confidence to the robustness of the observed differences. The moderate interaction strengths observed here are consistent with a dynamic equilibrium between monomeric and oligomeric receptor populations, as expected for GPCRs operating in the crowded and heterogeneous environment of the plasma membrane.

The two receptor variants differ in the architecture of their fluorescent protein fusions: MC4R-A contains a membrane anchor in helix VIII followed by a short flexible linker to the fluorescent protein, whereas MC4R-B2 lacks this anchor but features a longer intracellular extension preceding the tag. Because linker length, flexibility, and anchoring can in principle influence fluorophore separation and orientation, these constructs also serve as an internal methodological control for potential linker-dependent biases in FRET-based distance and interaction estimates. Notably, we did not observe systematic differences in apparent distances or FRET efficiencies between MC4R-A and MC4R-B2 that could be attributed to linker architecture. This suggests that, within the resolution and sensitivity of our measurements, linker composition did not measurably distort the inferred interaction parameters. Nevertheless, linker structure and fluorophore mobility remain important considerations for quantitative FRET studies. A systematic investigation of linker length and sequence effects across standardized constructs would be valuable for establishing broadly transferable design rules and minimizing the need for extensive construct-specific simulations in future studies (Basak et al. 2021). The current construct design with the C-terminal FP tagging limits the ability to differentiate competing structural models. Strategically placed FPs (for instance in the N-terminus, or an intra- or extracellular loop) may increase our ability to resolve distinct dimerization interfaces.

### 4.3 Challenges in live-cell FLIM and spectroscopic measurements

Accurate analysis is a central prerequisite for the correct interpretation of the acquired imaging data. Prior to this study, the lack of user-friendly, open-source analysis workflows presented a major obstacle. Many laboratories rely on closed vendor platforms or custom scripts written in MATLAB, which hinders reproducibility and accessibility. In response, our step-by-step described workflow provides a fully open-source, beginner-friendly toolkit composed of GUI-driven applications and Jupyter notebooks (Greife, et al. 2025). This approach not only lowers the barrier for new users but also maximizes transparency and reproducibility of PIE-FLIM data analysis. A further central prerequisite for quantitative FRET measurements is that the fluorescent protein (FP) fusion does not alter the biological behavior of the protein of interest. Additionally, tags must be positioned within a FRET-accessible range; *R_0_* values range ∼30–69 Å for common FP pairs listed in FPbase (Lambert 2019), thus giving an accessible range of up to 110 Å.

Moreover, accurate quantification of protein-protein interactions in living cells is inherently challenged by spatial heterogeneity in protein distribution, local concentration, and cellular context. In earlier FLIM studies regions of interest were selected manually, e.g. membrane-associated receptor populations or specific intracellular compartments (Kravets, et al. 2016). Such manual region selection can be subjective, difficult to reproduce, and scales poorly to large datasets. Here, we demonstrate an automated PIE-FLIM image segmentation analysis that can improve robustness and information yield by subdividing cells into biologically meaningful subregions. Our approach analyzes biomolecular states as a function of the local environment and protein density. Such spatial stratification reduces averaging, enhances sensitivity to intracellular heterogeneity, and expands the effective concentration range, central for the reliable determination of oligomerization constants. An integrated analysis workflow enables the incorporation of image- and spectroscopy-based features. Local intensity distributions, molecular density proxies, shape descriptors, and N&B–derived estimates can be coupled to fluorescence lifetime, anisotropy, and FRET efficiency measurements. Linking these observables allows improving the discrimination between monomeric, dimeric, and higher-order assemblies. By shifting from manually curated regions to reproducible, feature-based spatial analysis, segmentation becomes a key component of quantitative live-cell interaction studies rather than a purely technical preprocessing step.

## 5 Outlook

We demonstrated how to disentangle oligomerization states in living cells using homo- and heteroFRET. Nevertheless, live-cell spectroscopy remains challenging due to constraints in sample preparation and data analysis. High-quality fluorescence lifetime imaging requires careful microscope calibration, control of imaging conditions, and limiting phototoxicity/photobleaching due to high irradiances. Likewise, quantitative interpretation of FRET signals is complicated by the need to disentangle photophysics, labeling stoichiometry, dipole orientations, and receptor dynamics (Wallrabe et al. 2006). Advances in FP engineering - including brighter, photostable variants (e.g., mStayGold, mScarlet (Gadella et al. 2023, Goedhart and Gadella 2024) and red-shifted donor-acceptor combinations (Zhang et al. 2023) - have helped mitigate phototoxicity and improve live-cell compatibility by lowering the excitation density and required FP overexpression for detectability.

Still, the efficient use of the limited photon budget motivates approaches that maximize the information extracted from every photon by combining homo- and heteroFRET. Generally, homoFRET provides a unique advantage: it requires only a single fluorophore species, eliminating chromatic aberrations and freeing a second spectral channel for independent biological markers such as biosensors of cell state, second messengers, or receptor activity. This is especially valuable in live-cell imaging, where phototoxicity, acquisition speed, and multiplexing capacity are often limiting (Liput, et al. 2020, Wallrabe, et al. 2006). The combination of fluorescence anisotropy and fluorescence lifetime allows simultaneous visualization of homo- and heteroFRET in parallel when studying homotypic and heterotypic protein interactions (Liput, et al. 2020). Even though a joint framework for unified lifetime-anisotropy analysis that describes all acquired data is missing, we demonstrated consistent results for homo- and heteroFRET data acquired by pulsed-interleaved excitation (Muller, et al. 2005) with multiparameter polarization resolved detection (Weidtkamp-Peters, et al. 2009) which simultaneously records hetero- and homoFRET data. Such combination is particularly well suited in GPCRs to dissect also state-dependent interactions with downstream signaling proteins and identifying conformational coupling within receptor complexes (Harkes et al. 2021). Mapping anisotropy versus lifetime provides a two-dimensional representation that separates homoFRET-induced depolarization from heteroFRET-driven lifetime shortening. This integrated representation not only unifies both FRET readouts but also enables approximation of relative interaction contributions, extending analytical power beyond what homo- or heteroFRET alone can achieve.

Fluorescent proteins can only provide limited structural information. However, we demonstrated how limited information can be useful to falsify competing structural models by combining structural prediction tools such as AlphaFold (Abramson, et al. 2024) or RoseTTAFold (Baek et al. 2021) with integrative coarse-grained approaches for multi-scale simulations in the Integrative Modeling Platform (Russel, et al. 2012). Central to relate structures to spectroscopic FRET observables, particularly in fluorescent proteins, is the handling of linker regions and the orientation factor *κ*². A persistent challenge in quantitative FRET analysis is the uncertainty which reflects the relative orientation of donor and acceptor transition dipoles. For fluorescent proteins, rotational diffusion is slow relative to their fluorescence lifetime, rendering the commonly assumed isotropic average of *κ*² = 2/3 generally invalid (Hummer and Szabo 2017, Meer, et al. 2013, Peulen, et al. 2017). As a result, absolute distance and interaction estimates derived from FRET measurements may be systematically biased if dipole orientation constraints are not considered. We demonstrated how *κ*² distributions, the spatial freedom, steric constraints, and preferred orientations of fluorescent protein tags within the molecular assembly can be handled *in silico*. A direct experimental access to these distributions is limited, and brute-force molecular dynamics simulations remain computationally demanding for large, membrane-embedded protein–fluorophore systems. Thus, future iterative refinement between structural modeling and experimental fluorescence data offers a promising strategy to reduce uncertainty in *κ*² and to improve the quantitative interpretation of interaction parameters for calibrated experiments and simulations. Beyond improving affinity estimates, such integrative approaches also provide a structural framework for understanding receptor organization and conformational coupling within oligomeric assemblies.

## 7 Author Contributions

- AG - designed study, checked and wrote data analysis manuals, created figures and wrote manuscript
- RL - checked data analysis manuals, analyzed MC4R-A data, wrote manuscript
- PSK - performed cell culture and sample preparation, acquired data, analyzed MC4R-B2 data
- KGH - designed study, acquired funding, edited manuscript
- KH - designed study, developed data analysis workflow, wrote data analysis manuals, created figures and wrote manuscript
- TOP - designed study, developed data analysis workflow, wrote analysis software and manuscript

## 9 Acknowledgments

The Core Unit Fluorescence Imaging of the RVZ/JMU, especially Mike Friedrich for help with the LSM980, and Kerstin Jansen for help with cell culture.

## 11 Supplementary Material

- Supplementary Figures
- Example Data with Step-by-Step Data Export Manual, Lifetime Fits Analysis Manual, Gaussian Distance Fits Analysis Manual and Anisotropy Analysis Manual (10.5281/zenodo.17869433)
- Raw data files and cell masks (10.5281/zenodo.18175060)

## 11 Data and Code Availability Statement

The full data set (10.5281/zenodo.18175060) and an example data set including the scripts and step-by-step manuals to generate the results in this study have been uploaded to zenodo (10.5281/zenodo.17869433). *ChiSurf* (Peulen 2025) and *tttrlib* (Peulen, et al. 2025) are open-source software and can be downloaded from the website (https://www.peulen.xyz/downloads/chisurf/), and GitHub repository (https://github.com/Fluorescence-Tools/tttrlib).

## 12 Generative AI Statement

The authors declare that Generative AI was used in the editing of this manuscript to improve readability. Grammarly and ProWritingAid were used for English correction and editing.

