## Supplementary Figures and Tables for "Bound or unbound: Mapping and monitoring receptor oligomerization using time-resolved fluorescence"

### 1 Supplementary Figures and Tables

#### 1.1 Supplementary Figures

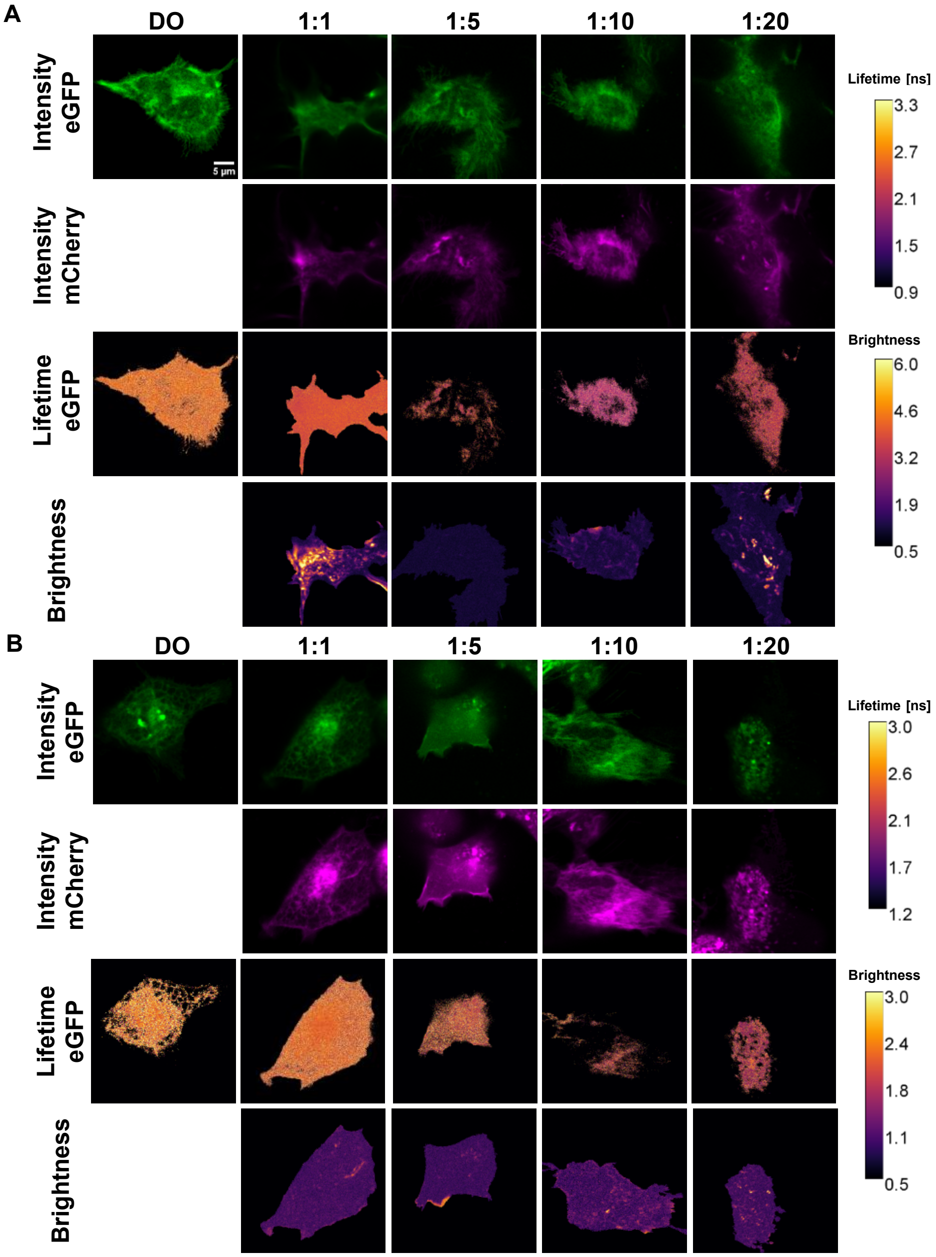

**Supplementary Figure 1. Fluorescence lifetime images of representative cells (A)** eGFP fluorescence intensity in the "prompt" time window (eGFP excitation, 485 nm), mCherry fluorescence intensity in the "delay" time window (mCherry excitation, 561 nm), mean lifetime and brightness images for selected MC4R-A cells either transfected with the MC4R-eGFP construct alone (DO) or in a 1:1, 1:5, 1:10 or 1:20 ratio of MC4R-A-eGFP and MC4R-A-mCherry. The total plasmid amount was kept constant. **(B)** Same as (A) for the MC4R-B2 construct. The mean lifetimes were computed for pixels with more than 15 photons.

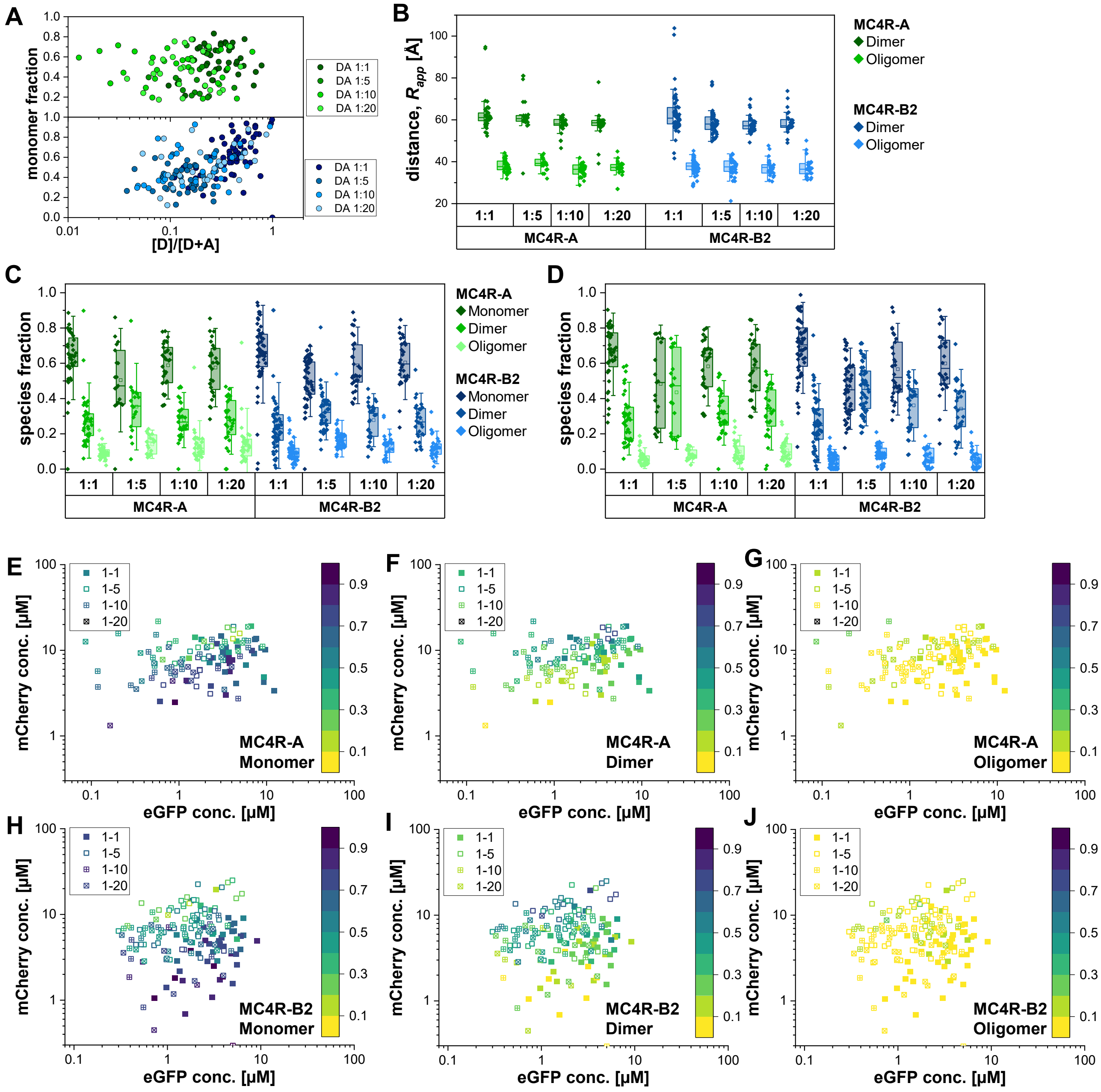

**Supplementary Figure 2. Concentration-dependent changes of MC4R monomer, dimer, and oligomer fractions. (A)** The monomer fractions (determined by a Gaussian model) depend on the donor to total protein concentration ratio ([D]/[D]+[A]) (MC4R-A, green; MC4R-B2, blue). **(B)** Fitted apparent mean inter-fluorophore distances for the two-Gaussian distance model. **(C)** Corresponding species fractions. **(D)** Species fraction for the **global** two-Gaussian distance model. (Dark color: monomer, mid color: dimer, lighter color: oligomer). **(E-G)** The observed species fractions depend on the donor (eGFP-tagged) and acceptor (mCherry-tagged) concentration. MC4R-A monomer (E), dimer (F), and oligomer (G) species fractions are color-coded (0 %, yellow; 100 %, dark blue). Filled, open, crossed, circles symbol corresponds to a 1:1, 1:5, 1:10, and 1:20 eGFP to mCherry transfection ratio. **(H-J)** Same as (E-G) for MC4R-B2.

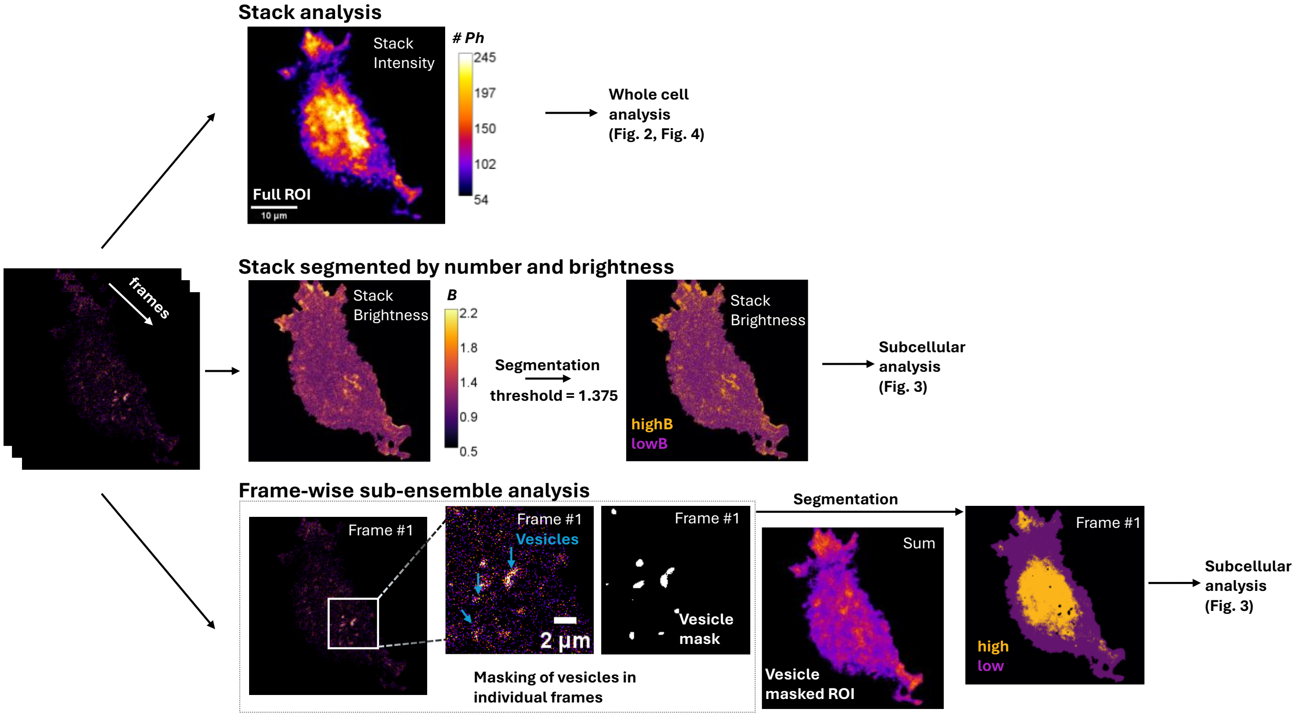

**Supplementary Figure 3. Subcellular analysis of the cell ROIs.** The FLIM time-series images were segmented and analyzed using three different approaches. **(Top)** In Figures 3 and 5, the whole cell was used as a region of interest (ROI). The cell outline was defined based on the stacked image. **(Mid)** In a 2$^{nd}$ approach, the stacked image was analyzed using the number and brightness approach, and the complete cell ROI was split into a low and high brightness region using a threshold of $B$ =1.375. The threshold was set based on a manual inspection of multiple datasets. **(Bottom)** In the 3$^{rd}$ approach, high-intensity vesicles were detected on a frame-by-frame basis and removed from the cell ROI. The remaining fluorescence intensity was split frame-by-frame in a low and high intensity region using Otsu's thresholding.

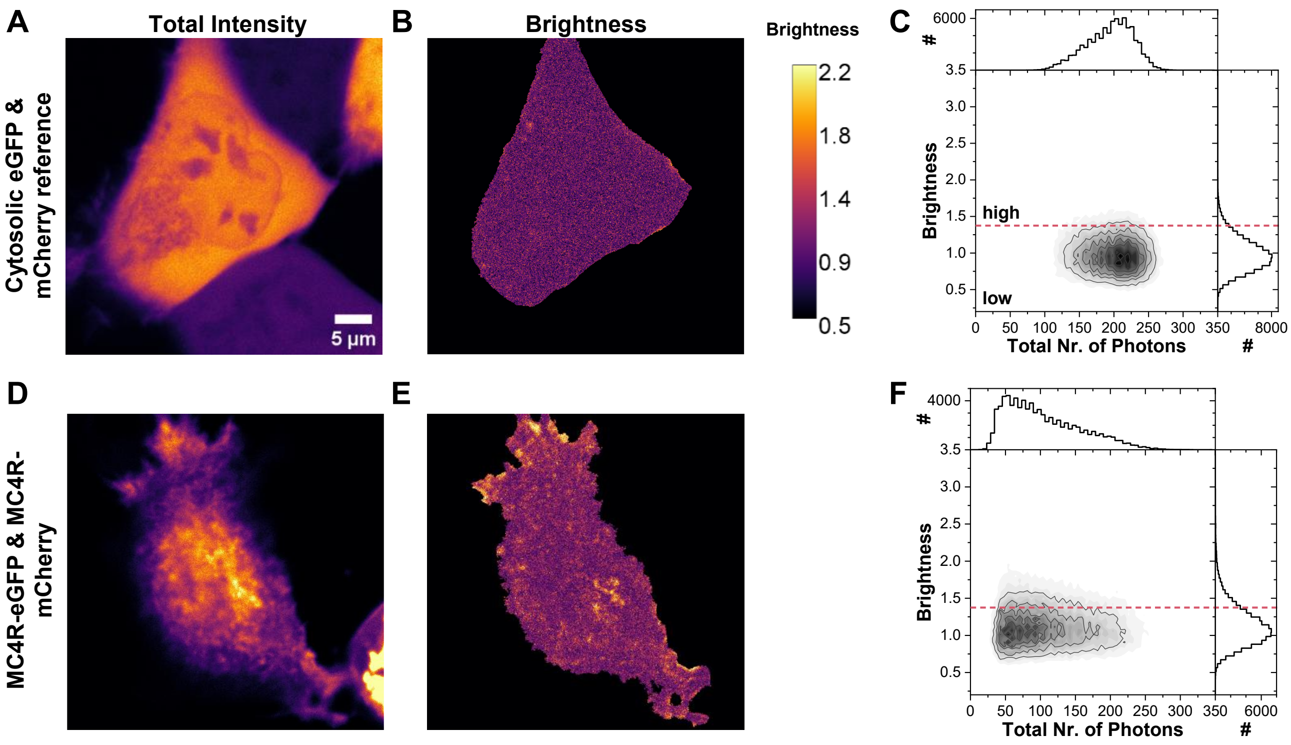

**Supplementary Figure 4. Number & Brightness approach. (A)** Intensity image of a cell co-transfected in a 1:1 ratio with cytosolic, monomeric eGFP and mCherry. **(B)** Brightness image of the same cell calculated based on the intensity fluctuations of all channels and time windows (green prompt, red prompt and red delay). **(C)** Pixel-wise 2D histogram of total intensity vs brightness. **(D)-(F)** Same as (A-C) for a cell co-transfected with MC4R-A-eGFP and MC4R-A-mCherry. Scale bar 5 µm.

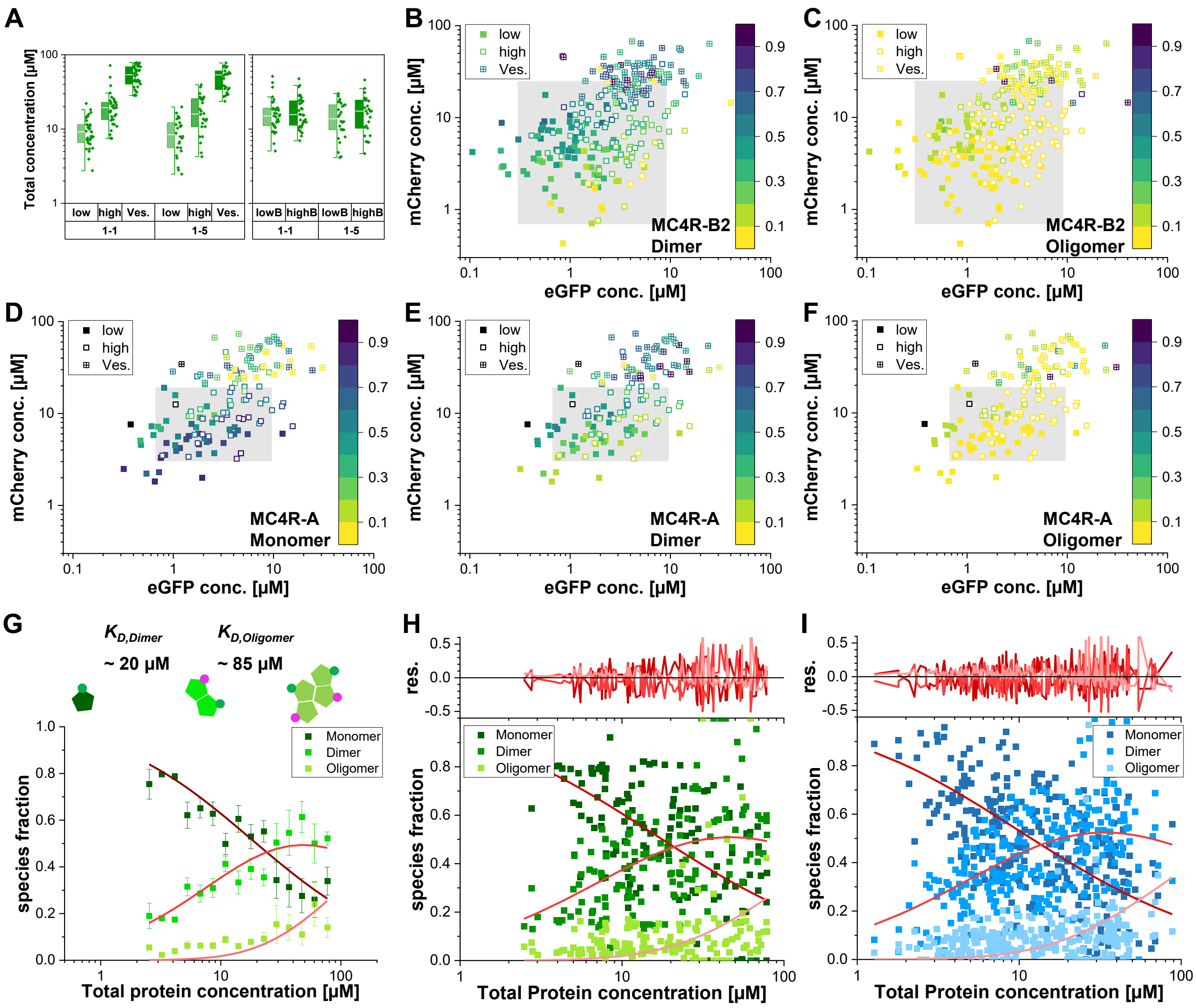

**Supplementary Figure 5. Fit results for the intensity-based sub-segmentation for MC4R-A and MC4R-B2. (A)** Boxplot of the total protein concentration for MC4R-A showing the accessible protein concentration distribution in the individual sub-segmented regions. **(B, C)** MC4R-B2-eGFP vs MC4R-B2-mCherry concentration color-coded by dimer (B) or oligomer (C) fraction obtained from fitting the intensity-based sub-segmentation globally with the fixed dimer and oligomer distance. Species fractions are color-coded (10 % (yellow) - 90 % (dark blue)). The shaded area shows the accessible concentration range of the 1:1 and 1:5 transfection ratio samples without the sub-segmentation. **(D) - (F)** Color-coded monomer (D), dimer (E), and oligomer (F) fractions for MC4R-A from fitting the intensity-based sub-segmentation globally with the fixed dimer and oligomer distance. **(G)** Average monomer (dark green), dimer (green), and oligomer (light green) fraction vs. total protein concentrations. Species fractions were fitted with the oligomerization model (dark red, red, and light red). Error bars show the standard error of the mean. **(H)** Species fractions of the global fit of MC4R-A were fitted with the Oligomerization model (**Eqs. 17-19**). Color-code: Dark green/red: Monomer, green/red: dimer, light green/red: oligomer. **(I)** Same as (H) for MC4R-B2 (Dark blue/red: Monomer, blue/red: dimer, light blue/red: oligomer). The fit results for (H, I) are summarized in **Supplementary Table 3.**

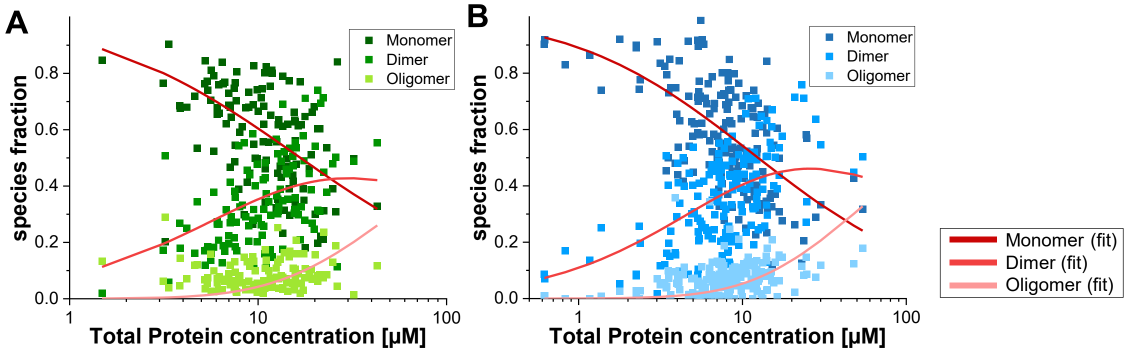

**Supplementary Figure 6. Oligomerization model for MC4R-A and MC4R-B2 full cell ROIs. (A, B)** Species fractions of the global two-Gaussian distances model of MC4R-A (A) and MC4R-B2 (B) of the full ROIs were fitted with the Oligomerization model (**Eqs. 17-19**). The fit results are summarized in **Supplementary Table 3**.

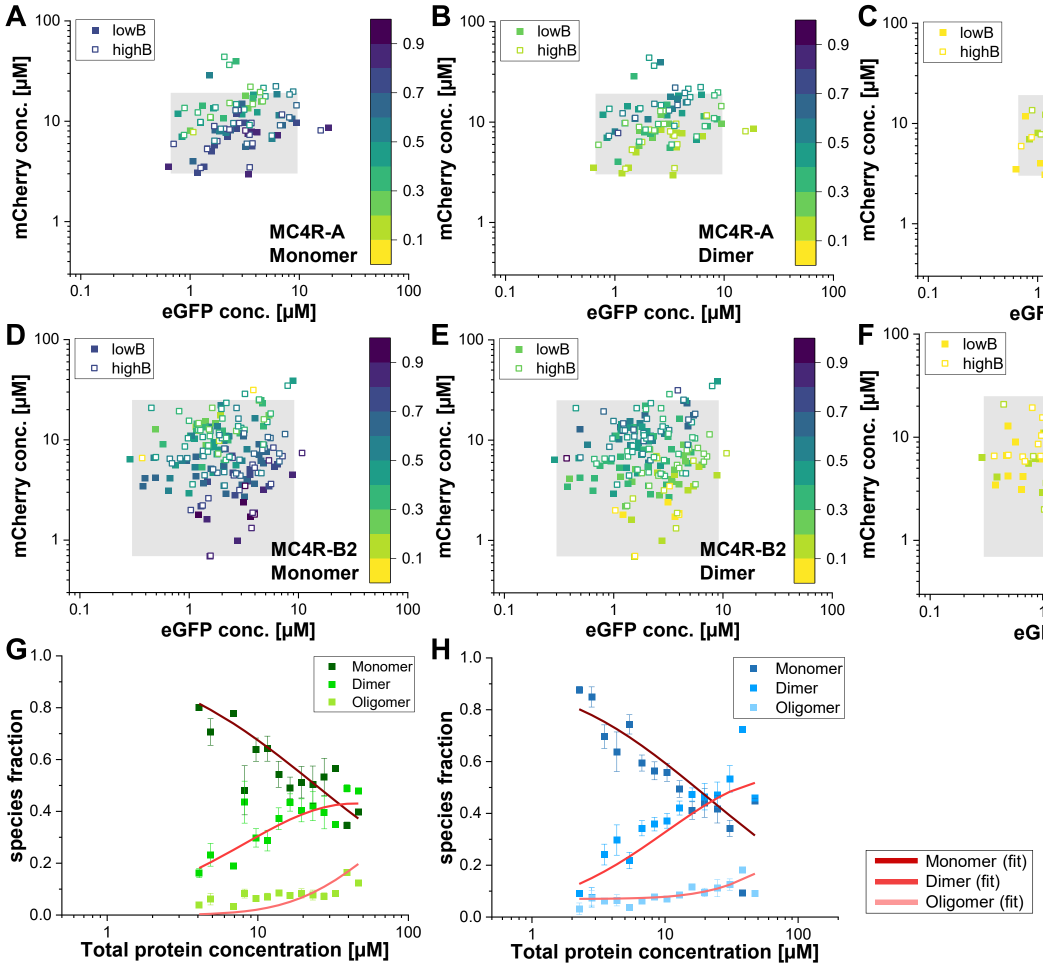

**Supplementary Figure 7. Results of the Oligomerization model for the MC4R-A and MC4R-B2 brightness-based sub-segmentation. (A)-(C)** MC4R-A-eGFP vs MC4R-A-mCherry concentrations color-coded for monomer (A), dimer (B) or oligomer (C) fractions obtained from fitting the brightness-based sub-segmentation globally with the fixed dimer and oligomer distance. Species fractions are color coded (10 % (yellow) - 90 % (dark blue)). The shaded area shows the accessible concentration range of the 1:1 and 1:5 transfection ratio samples without sub-segmentation. **(D)-(F)** Same as (A)-(C) for MC4R-B2. **(G)** Average monomer (dark green), dimer (green) and oligomer (light green) fractions vs the total protein concentrations obtained from the brightness-based sub-segmentation. Species fractions were fit with the Oligomerization model (**Eqs. 17-19**). Color-code: dark red, red and light red. Error bars are standard errors of the mean. **(H)** Same as (G) for MC4R-B2 (Dark blue/red: Monomer, blue/red: dimer, light blue/red: oligomer). The fit results for (G, H) are summarized in **Supplementary Table 2**. Note that $K_{D,Oligomer}$ is higher than the assessed concentration range.

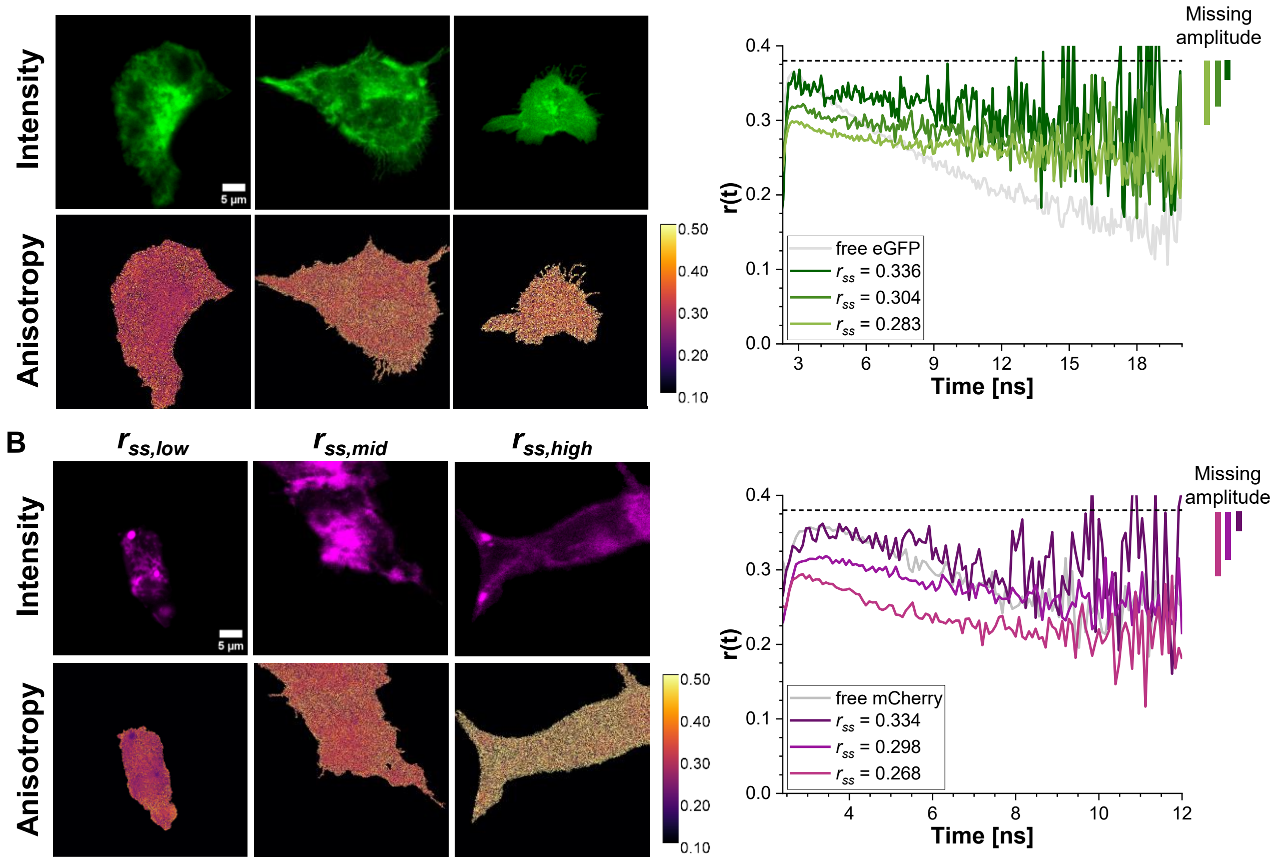

**Supplementary Figure 8. Steady-state fluorescence anisotropy images of selected cells. (A)** Green intensity in the "prompt" time window and steady-state anisotropy of three cells with overall low, mid and high steady-state anisotropy of cells transfected with MC4R-A-eGFP (DO). The time-resolved anisotropy is shown on the side. **(B)** Acceptor intensity in the "delay" time window and steady-state anisotropy of three cells with overall low, mid and high steady-state anisotropy of cells transfected with MC4R-B2-mCherry. The time-resolved anisotropy is shown on the side. For the green channels a g-factor of 0.946 and for the red channels a g-factor of 0.995 was used. The data were exported with a time resolution of 80 ps/bin to reduce data noise. Scale bar: 5 µm.

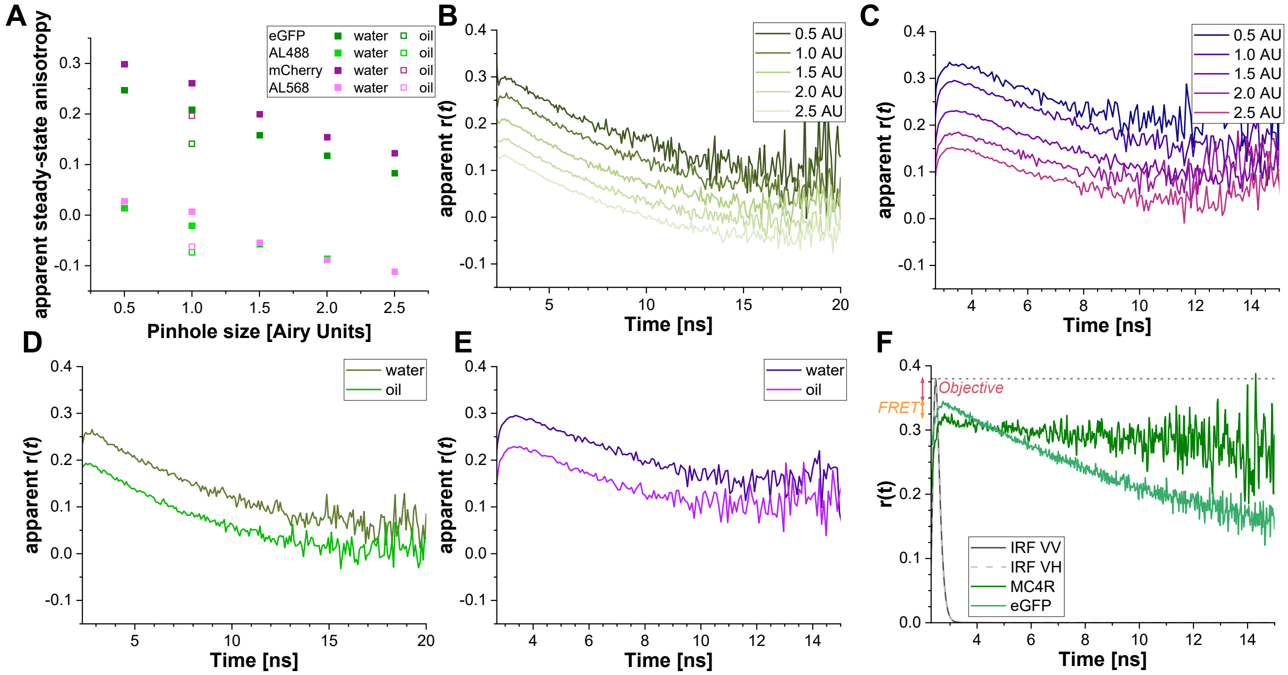

**Supplementary Figure 9. Characterization of the depolarization through the microscope objective. (A)** Uncorrected, apparent steady-state anisotropy of eGFP, Alexa488, mCherry and Alexa568 solutions determined for our 40x water objective (NA 1.2) or the 63x oil objective (NA1.4) and different pinhole sizes (0.5 - 2.5 Airy Units, AU). **(B)** Time-resolved anisotropies of eGFP. **(C)** Time-resolved anisotropies of mCherry. **(D)** Comparison of the apparent time-resolved anisotropy of eGFP measured with the 40x water objective (NA 1.2) and the 63x oil objective (NA 1.4). **(E)** Same as (D) for mCherry. The apparent anisotropy in panels (A) – (E) is uncorrected for the g-factor and was calculated as $r_{ss,app} = \frac{\sum I_{VV}(t) - \sum I_{VH}(t)}{\sum I_{VV}(t) + 2 * \sum I_{VH}(t)}$ and $r_{app}(t) = \frac{I_{VV}(t) - I_{VH}(t)}{I_{VV}(t) + 2 * I_{VH}(t)}$. To reduce data noise, the data was exported with a time resolution of 80 ps/bin. **(F)** For a fluorophore with a fundamental anisotropy $r_0 = 0.38$, the time-resolved anisotropy r(*t*) should initially be 0.38. Missing amplitudes may be due to (*i*) depolarization of the objective or (*ii*) very fast homoFRET processes. Comparing a reference dye or sample (here, an eGFP solution), in which homoFRET processes can be excluded, provides information on whether homoFRET occurs in the sample of interest (here, an MC4R measurement). For illustration, the IRF is also shown, and its height was normalized to 0.38.

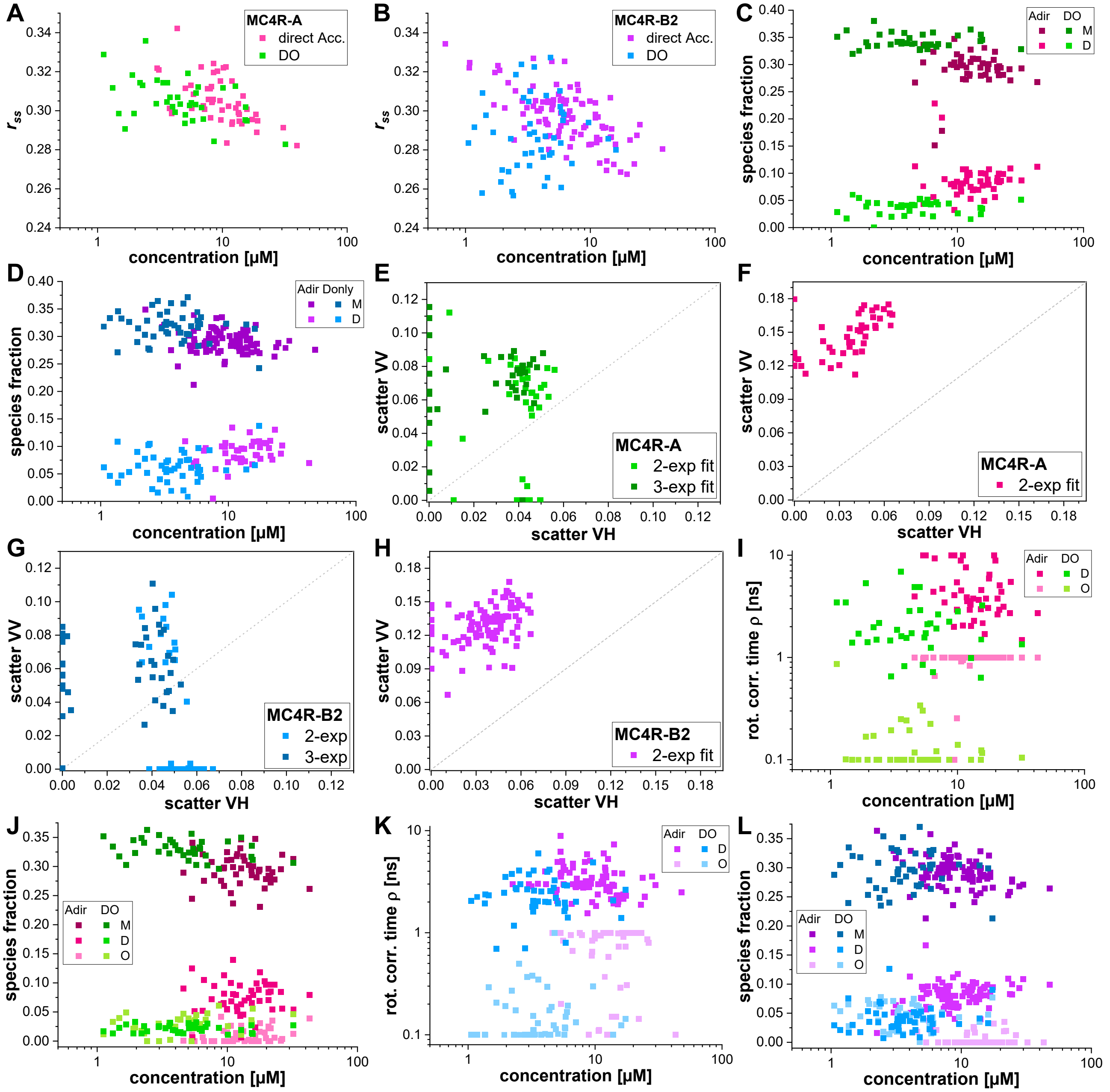

**Supplementary Figure 10. Comparison of dimer and oligomer models in time-resolved anisotropy measurements. (A)** Concentration-dependence of the steady-state anisotropy of DO (green) and directly excited acceptor (magenta) of the 1:1 and 1:5 co-transfected MC4R-A samples. **(B)** Same as (A) for MC4R-B2 (DO: blue, directly excited acceptor: violet). **(C)** Species fractions obtained from the dimer model for MC4R-A with a free dimer relaxation time but a fixed rotational correlation time. The concentration axis reflects the total protein concentration. **(D)** Same as (C) for MC4R-B2 (D: Dimer, M: Monomer). **(E, F)** Scatter obtained from in the dimer model (light green, DO (E); directly excited acceptor (F)) or the oligomer model (dark green, DO, (E)) in the parallel (VV) and perpendicular (VH) detection channels for MC4R-A. **(G, H)** Same as (E-F) for MC4R-B2. **(I, J)** Relaxation times (I) and species fractions (J) from the oligomer model with freely floating Oligomer and Dimer relaxation times (fixed molecule rotation) for MC4R-A (D: Dimer, M: Monomer, O: Oligomer). **(K, L)** Same as (I-J) for MC4R-B2.

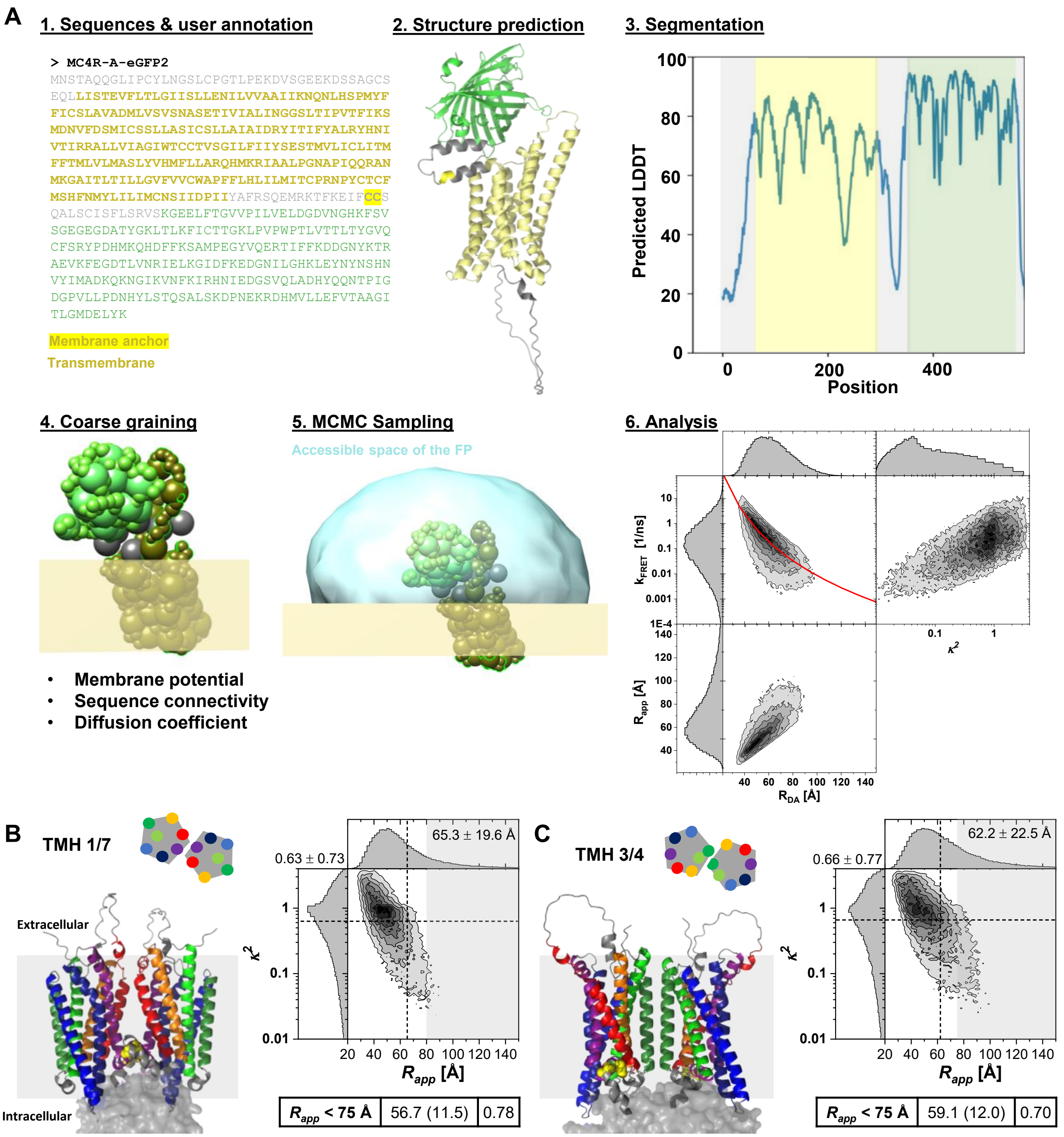

**Supplementary Figure 11. Simulation workflow to estimate the spatial fluorescent proteins distribution on target proteins and resulting FRET efficiencies for protein complexes.** (**A**) The fluorescent protein simulation implemented in FPSIMP (https://github.com/fluorescence-tools/fpsimp) follows a 6-step procedure: **(1)** In a first step the complete sequence of the protein of interest and the fluorescent protein is entered and annotated, e.g. transmembrane regions or membrane anchors are defined. **(2)** The structure of the tagged proteins is predicted using AlphaFold3. (3) Rigid elements (helices, β-sheets) and flexible regions are identified based on their predicted LDDT (local distance difference test), which describes the per-residue local confidence of the generated structure. **(4)** The integrative modeling platform (IMP) is used to add information such as membrane potential (i.e. such that the membrane anchor residues will stay close to/in the membrane) and to create a coarse-grained model. **(5)** The coarse-grained model is used in Markov-Chain Monte-Carlo (MCMC) sampling to determine the accessible space of the fluorescent protein (FP, light blue halo). Here, > 10 000 structures / orientations are generated. **(6)** Using the position and the orientation of the chromophore in eGFP and mCherry, the inter-fluorophore distance distribution, $R_{DA}$, and the orientation factor distribution, $\kappa^2$, can be obtained. $R_{app}$ describes the distance obtained from fitting. The red line indicates the observed distance for $\kappa^2 = 2/3$. (**B, C**) AlphaFold3 predicts different dimerization interfaces for MC4R-A. Simulated apparent inter-fluorophore distances for the TMH 1/7 (B) and the TMH 3/4 (C) dimerization model. The table indicates the distribution mean and width (in brackets) when assigning all models with $R_{app}$ > 75 Å to $x_{noFRET}$ (22-30%). The color code of helices is identical to **Figure 6D** in the main text. eGFP and mCherry are shown as semitransparent surface and the gray shaded area indicates the cell membrane. Dashed lines: Mean values for $\kappa^2$ and $R_{app}$, numbers in the 2D histogram report mean and width of the distributions.

#### 1.2 Supplementary Tables

**Supplementary Table 1.** Amino acid sequences for the MC4R-A and MC4R-B2 fluorescent protein tagged constructs. The eGFP (green) and mCherry (magenta) sequences are highlighted. Orange residues mark the membrane anchor in MC4R-A, bold sequences mark the intracellular helix VIII andflexible intracellular residues.

| ***Construct*** | ***Sequence*** |
|---|---|
| MC4R-A-eGFP | MNSTAQQGLI PCYLNGSLCP GTLPEKDVSG EEKDSSAGCS EQLLISTEVF<br>LTLGIISLLE NILVVAAIIK NQNLHSPMYF FICSLAVADM LVSVSNASET<br>IVIALINGGS LTIPVTFIKS MDNVFDSMIC SSLLASICSL LAIAIDRYIT<br>IFYALRYHNI VTIRRALLVI AGIWTCCTVS GILFIIYSES TMVLICLITM<br>FFTMLVLMAS LYVHMFLLAR QHMKRIAALP GNAPIQQRAN MKGAITLTIL<br>LGVFVVCWAP FFLHLILMIT CPRNPYCTCF MSHFNMYLIL IMCNSIIDPI<br>I**YAFRSQEMR KTFKEIFCCS QALSCISFLS RVS**KGEELFT GVVPILVELD<br>GDVNGHKFSV SGEGEGDATY GKLTLKFICT TGKLPVPWPT LVTTLTYGVQ<br>CFSRYPDHMK QHDFFKSAMP EGYVQERTIF FKDDGNYKTR AEVKFEGDTL<br>VNRIELKGID FKEDGNILGH KLEYNYNSHN VYIMADKQKN GIKVNFKIRH<br>NIEDGSVQLA DHYQQNTPIG DGPVLLPDNH YLSTQSALSK DPNEKRDHMV<br>LLEFVTAAGI TLGMDELYK |
| MC4R-A-mCherry | MNSTAQQGLI PCYLNGSLCP GTLPEKDVSG EEKDSSAGCS EQLLISTEVF<br>LTLGIISLLE NILVVAAIIK NQNLHSPMYF FICSLAVADM LVSVSNASET<br>IVIALINGGS LTIPVTFIKS MDNVFDSMIC SSLLASICSL LAIAIDRYIT<br>IFYALRYHNI VTIRRALLVI AGIWTCCTVS GILFIIYSES TMVLICLITM<br>FFTMLVLMAS LYVHMFLLAR QHMKRIAALP GNAPIQQRAN MKGAITLTIL<br>LGVFVVCWAP FFLHLILMIT CPRNPYCTCF MSHFNMYLIL IMCNSIIDPI<br>I**YAFRSQEMR KTFKEIFCCS QALSCISFLS RVS**KGEEDNM AIIKEFMRFK<br>VHMEGSVNGH EFEIEGEGEG RPYEGTQTAK LKVTKGGPLP FAWDILSPQF<br>MYGSKAYVKH PADIPDYLKL SFPEGFKWER VMNFEDGGVV TVTQDSSLQD<br>GEFIYKVKLR GTNFPSDGPV MQKKTMGWEA SSERMYPEDG ALKGEIKQRL<br>KLKDGGHYDA EVKTTYKAKK PVQLPGAYNV NIKLDITSHN EDYTIVEQYE<br>RAEGRHSTGG MDELYK |
| MC4R-B2-eGFP | MNSTAQQGLI PCYLNGSLCP GTLPEKDVSG EEKDSSAGCS EQLLISTEVF<br>LTLGIISLLE NILVVAAIIK NKNLHSPKYF FICSLAVADM LVSVSNASET<br>IVIALFNGGS LTIPVTFIKS MDNVFNSMIC SSLLASICSL LAIAIDRYIT<br>IFYALRYHDI VTIRRALLVI GSIWTCCTVS GILFIIYSES TVVLICLITM<br>SFTVLVLMAS LYVHMFLLAR QHMKRIGALP GNAAIQQRAN MKGAITLTIL<br>LGVFVFCWAP FFLHLILMIT CPRNPYCTCF MSHFNMYLIL IMCNSVIDPI<br>I**YAFRSQEMR KTFKKIFSQA RVLAFCETLQ VVHLSRVS**KG EELFTGVVPI<br>LVELDGDVNG HKFSVSGEGE GDATYGKLTL KFICTTGKLP VPWPTLVTTL<br>TYGVQCFSRY PDHMKQHDFF KSAMPEGYVQ ERTIFFKDDG NYKTRAEVKF<br>EGDTLVNRIE LKGIDFKEDG NILGHKLEYN YNSHNVYIMA DKQKNGIKVN<br>FKIRHNIEDG SVQLADHYQQ NTPIGDGPVL LPDNHYLSTQ SALSKDPNEK<br>RDHMVLLEFV TAAGITLGMD ELYK |
| MC4R-B2-mCherry | MNSTAQQGLI PCYLNGSLCP GTLPEKDVSG EEKDSSAGCS EQLLISTEVF<br>LTLGIISLLE NILVVAAIIK NKNLHSPKYF FICSLAVADM LVSVSNASET<br>IVIALFNGGS LTIPVTFIKS MDNVFNSMIC SSLLASICSL LAIAIDRYIT<br>IFYALRYHDI VTIRRALLVI GSIWTCCTVS GILFIIYSES TVVLICLITM<br>SFTVLVLMAS LYVHMFLLAR QHMKRIGALP GNAAIQQRAN MKGAITLTIL<br>LGVFVFCWAP FFLHLILMIT CPRNPYCTCF MSHFNMYLIL IMCNSVIDPI<br>I**YAFRSQEMR KTFKKIFSQA RVLAFCETLQ VVHLSRVS**KG EEDNMAIIKE<br>FMRFKVHMEG SVNGHEFEIE GEGEGRPYEG TQTAKLKVTK GGPLPFAWDI<br>LSPQFMYGSK AYVKHPADIP DYLKLSFPEG FKWERVMNFE DGGVVTVTQD<br>SSLQDGEFIY KVKLRGTNFP SDGPVMQKKT MGWEASSERM YPEDGALKGE<br>IKQRLKLKDG GHYDAEVKTT YKAKKPVQLP GAYNVNIKLD ITSHNEDYTI<br>VEQYERAEGR HSTGGMDELY K |

**Supplementary Table 2.** The binned species fractions from the global two-Gaussian distance model for full ROIs and the intensity- and brightness-based sub-segmentation of MC4R-A and MC4R-B2 were fitted with the Oligomerization model. 95% confidence intervals (CI) were determined by 500 rounds of bootstrapping. [] indicates CI.

| ***Parameter*** | ***MC4R-A (full ROI)*** | ***MC4R-A Intensity*** | ***MC4R-A Brightness*** | ***MC4R-B2 (full ROI)*** | ***MC4R-B2 Intensity*** | ***MC4R-B2 Brightness*** |
|---|---|---|---|---|---|---|
| Used data | 1-1, 1-5, 1-10, 1-20 | 1-1, 1-5 | 1-1, 1-5 | 1-1, 1-5, 1-10, 1-20 | 1-1, 1-5 | 1-1, 1-5 |
| $n_{points}$ | 39 | 45 | 42 | 39 | 45 | 42 |
| $K_{D,Dimer}$ [µM] | 20.9 [17.1-25.3] | 19.9 [16.2-23.9] | 25.7 [19.7 – 32.0] | 15.5 [12.6 – 18.5] | 15.6 [12.7 – 21.5] | 17.4 [14.6 – 21.4] |
| $K_{D,Oligomer}$ [µM] | 69.2 [22.8 - 122] | 85.9 [43.7 – 127] | 54.0 [36.0 – 71.1] | 57.2 [16.7 – 77.1] | 81.7 [41.6 – 112] | 57.0 [26.3 – 89.7] |

**Supplementary Table 3.** The species fractions from the global two-Gaussian distance model for full ROIs and the intensity- and brightness-based sub-segmentation of MC4R-A and MC4R-B2 were fitted with the Oligomerization model. 95% confidence intervals (CI) determined by 500 rounds of bootstrapping. [] indicates CI.

| ***Parameter*** | ***MC4R-A (full ROI)*** | ***MC4R-A Intensity*** | ***MC4R-A Brightness*** | ***MC4R-B2 (full ROI)*** | ***MC4R-B2 Intensity*** | ***MC4R-B2 Brightness*** |
|---|---|---|---|---|---|---|
| Used data | 1-1, 1-5, 1-10, 1-20 | 1-1, 1-5 | 1-1, 1-5 | 1-1, 1-5, 1-10, 1-20 | 1-1, 1-5 | 1-1, 1-5 |
| $n_{points}$ | 348 | 429 | 288 | 513 | 819 | 558 |
| $K_{D,Dimer}$ [µM] | 20.7 [21.0 – 28.9] | 19.9 [16.4 – 24.0] | 24.6 [19.8 – 29.7] | 14.6 [12.8 – 17.0] | 13.0 [11.5 – 14.9] | 15.6 [C13.7 – 18.1] |
| $K_{D,Oligomer}$ [µM] | 29.7 [26.8 – 42.6] | 77.1 [41.5 – 119.0] | 39.5 [32.2 – 46.0] | 31.4 [23.3 – 41.8] | 59.6 [41.8 – 83.6] | 34.4 [27.9 – 42.1] |

**Supplementary Table 4.** The species fractions obtained from the time-resolved anisotropy analysis of MC4R-A and MC4R-B2 were fitted with the Dimerization model. CI = 95% confidence interval as determined by 500 rounds of bootstrapping. [] indicates CI.

| ***Parameter*** | ***MC4R-A (Anisotropy)*** | ***MC4R-B2 (Anisotropy)*** |
|---|---|---|
| Used data | 1-1, 1-5, 1-10, 1-20, DO | 1-1, 1-5, 1-10, 1-20, DO |
| $n_{points}$ | 170 | 298 |
| $K_{D,Dimer}$ [µM] | 56.0 [49.5 – 63.6] | 31.4 [28.6 – 34.2] |
